# A stochastic model of T cell expansion in activating micro-rod scaffolds and its continuum limit: Importance of IL-2 loading and scaffold homogeneity

**DOI:** 10.1101/2025.07.12.664022

**Authors:** M. S. Lacy, A. L. Jenner, P. R. Buenzli

## Abstract

T cells are immune cells that are known to be effective at killing cancer cells, however, an individual patient’s tumour-specific T cell counts are often insufficient to control cancer growths. Adoptive T cell therapy aims to address this by activating and expanding highly effective T cells *ex vivo* before injecting them into the patient to employ their cancer-killing functions. Recent experimental setups using activating micro-rod scaffolds have significantly improved T cell expansion over conventional methods, but there is still much to understand regarding the factors that maximise the expansion of functional T cells in these scaffolds. We present a stochastic agent-based model of T cell expansion alongside its continuum limit to simulate the average interactions between T cells and micro-rods, which enable us to explore several behaviours of the experimental system. Stochastic simulations demonstrate that T cell expansion is driven by activated cell clusters around micro-rods. Using our spatial models and a mean-field approximation, we discover that this cluster-driven expansion is most supported by scaffolds with initially homogeneous micro-rod concentrations. Our simulations also reveal that loading the T cell growth factor, interleukin-2 (IL-2), into micro-rod pores for secretion significantly prolongs expansion compared to the more conventional method of IL-2 supplementation.

## 1 Introduction

The immune system provides a sophisticated set of mechanisms as the body’s first line of defense that is able to fight against a wide variety of diseases, including cancer [1–3]. A key member of the human immune system is the effector T cell. Through antigen-presentation, effector T cells become activated and are able to induce death in cancer cells [1, 3, 4]. Unfortunately, cancer cells often rapidly develop the ability to evade T cells and diminish their therapeutic effectiveness [3, 5, 6]. Immunotherapies, such as checkpoint blockade and cancer vaccines, have been developed to help the immune system overcome cancer [2, 7, 8]. In this work we focus on adoptive T cell therapy (ACT), which is a cell-based cancer therapy that uses a patient’s own T cells initially derived from the tumour micro-environment. These cells are enhanced and expanded *ex vivo* (in a culture dish) before being infused back into the patient as activated anti-tumour effector T cells [9–11]. ACT has been used to successfully treat advanced melanoma, however, more research is required to improve the efficacy of ACT for other cancers and the efficiency of T cell expansion overall [9, 10, 12].

T cells are expanded *ex vivo* by mimicking the way that they are activated in the body [4, 10]. Normally, T cells receive co-stimulation from antigen-presenting cells (APCs) which display small pieces of the appropriate cancer antigen, known as peptides, on its surface [1, 3]. The T cell receptor (TCR) may be stimulated by a peptide-loaded major histocompatibility complex (pMHC) expressed on an APC (see Fig. 1a), which indicates to the T cell that the immune system has detected the presence of cancer. For successful activation, a secondary signal must be received through the T cell’s CD28 receptor, which occurs when it detects an appropriate molecule expressed on the APC, which we denote *α*CD28 [1, 3, 13]. T cells that receive this co-stimulation a sufficient number times reach a high level of activation, which improves the robustness of the cell’s response to cancer [14, 15]. After becoming activated, T cells begin to proliferate rapidly if they are exposed to the T cell growth factor, interleukin-2 (IL-2), which is a small signalling protein released by helper T cells and other immune cells [3,16]. This detection is driven by IL-2 receptors (IL-2R) which are more prominently expressed on the surface of T cells after activation [16] (see Fig. 1a).

**Figure 1.**
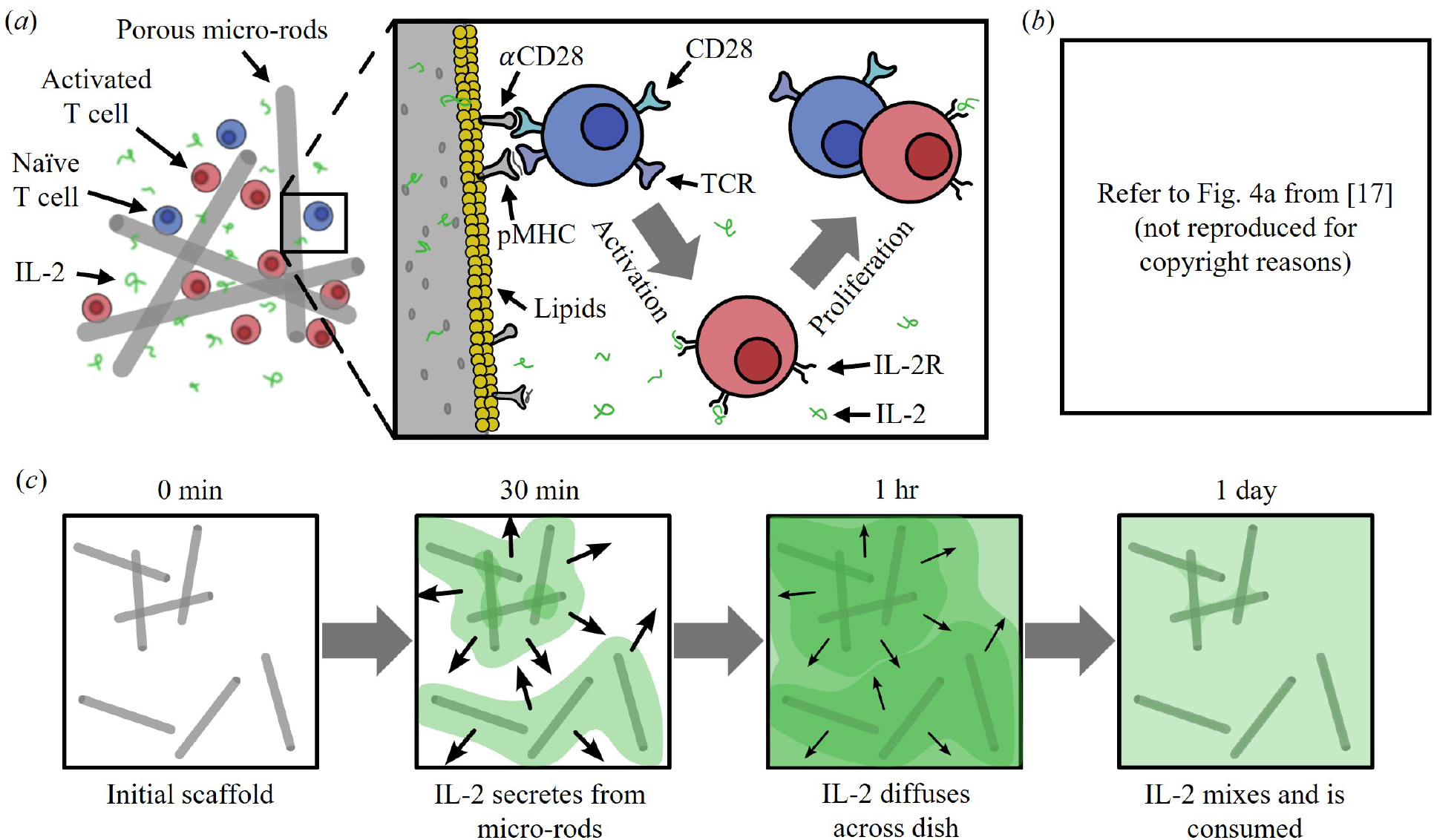
Summary of experimental micro-rod scaffold system. *(a)* Diagram of the interactions between T cells and micro-rods, detailing the activation of naïve T cells (blue) via co-stimulation by molecules (*α*CD28 and pMHC) attached to lipids on micro-rods. Proliferation occurs when activated T cells (red) detect the presence of IL-2 (green) using IL-2 receptors (IL-2R). *(b)* Representative bright-field micrograph of a micro-rod scaffold before co-culture with T cells (400 µm scale bar), reproduced from [17]. *(c)* Diagram of the sustained release of free IL-2 from micro-rods and their diffusion over time.

ACT requires the activation of T cells in culture dishes by utilising co-stimulatory signals similarly to those expressed by antigen-presenting cells [9]. In some experiments, biomaterials are used to imitate the activating functionality of APCs. For example, Dynabeads magnetic beads are artificial APCs which are commercially available and promote up to 1,000-fold expansion in T cells. To support T cell proliferation, scientists using these magnetic beads supplement the culture media with IL-2 [18–21]. As an alternative, Cheung *et al*. (2018) introduce a novel and exciting method that expands T cells using biomaterial scaffolds formed by APC-mimetic micro-rods (see Fig. 1), which provides much greater expansion than more conventional platforms such as Dynabeads magnetic beads [13, 17]. These micro-rods are surrounded by supported lipid bilayers, which mimic a cell’s surface and allows the attachment of activating stimuli such as pMHC and *α*CD28. The micro-rods are also mesoporous, which supports the loading of IL-2 inside micro-rods for sustained release during co-culture with T cells, imitating paracrine signalling in the immune system [13]. During T cell expansion, micro-rods form scaffolds in which T cells are able to freely move (Fig. 1b), promoting effective activation and proliferation. Unlike conventional methods, many aspects of the micro-rods are tunable, which allows for alterations to the T cell yield and composition of the resulting T cell population, such as the ratios of effector-to-helper and functional-to-dysfunctional T cells [13, 17]. This allows for appropriate adjustments to T cell expansion to better suit individual patients. However, the factors which drive and potentially optimise the expansion of functional T cells require further research.

Mathematical modelling is a powerful tool for conducting *in silico* (computer simulation) experiments and predicting improvements to laboratory experiments that are expensive or time-consuming to conduct. Existing ordinary differential equation (ODE) models of T cell expansion *in vivo* (in a living body) or *ex vivo* are able to predict the population growth of T cells in the presence of APCs and IL-2, however, they do not explicitly represent spatial dynamics and they do not consider expansion in the presence of tunable artificial APCs [22–25]. Existing partial differential equation (PDE) models and stochastic agent-based models (ABMs) provide frame-works for modelling T cell movement and spatial interactions with other entities [26–31]. To our knowledge, none are applied to investigate *ex vivo* T cell expansion for ACT. In this work, we present a novel spatial mathematical model of T cell expansion in activating micro-rod scaffolds. We predict ways in which expansion may be improved for more effective applications of ACT using micro-rod scaffolds. We also investigate some experimental observations described by Cheung *et al*., such as the effects of IL-2 loading and T cell clustering, with the goal of advancing our understanding of the mechanics which drive these observations. Our computational frameworks provide us with means to explore improvements to the design of activating micro-rods to maximise T cell expansion while maintaining their cancer-killing efficacy.

We consider a hybrid modelling framework, where T cells are described by a stochastic agent-based model in which individual T cells interact with activating micro-rods, and where IL-2 is modelled deterministically. We obtain the corresponding systems of partial and ordinary differential equations that approximate the stochastic model by taking its continuum limit. We simulate the ABM to investigate the behaviours of individual T cells, and compare the average stochastic predictions to the PDE model. We then compare our spatial models to the non-spatial ODE model, and investigate T cell expansion under various experimental designs.

## 2 Mathematical model

### 2.1 Agent-based model

Experimentally, T cells are expanded in quasi-two-dimensional wells or dishes where the culture media is more distributed in the horizontal plane than it is vertically. As such, we develop a two-dimensional (2D) model, where T cells, micro-rods and IL-2 occupy sites of an *L* × *L* square lattice, where *L* = 430 µm. The width of a lattice site is chosen to be *ε* = 5 µm (see Figs 2a,b) such that the centre of lattice site (*i, j*) lies at coordinates (*iε, j ε*), with *i, j* = 0, 1, 2, …, *R* where *R* = 86. The whole lattice represents a horizontal slice of the culture media (see Fig. 2a). We consider this slice to have a height of *h* = 5 µm, such that each lattice site contains all entities (cells, micro-rods and IL-2) within a small elementary volume of *ε* × *ε* × *h* = 125 µm^2^. We assume that any number of T cells or micro-rods may occupy one lattice site within the vertical constraints of the slice. We also assume there is no significant transport or heterogeneity in the vertical direction so that only one horizontal slice of the culture media is modelled. By considering a 2D domain, we perform conversions of some three-dimensional quantities to their analogues in the 2D simulations. These conversions are detailed in Sect. 2.5. For simplicity, micro-rods are assumed to be stationary, with discrete mass density *ρ*_*i, j*_ (in µg/µm^2^). Refer to Table 1 for a list of parameter values used in this model.

**Table 1.**
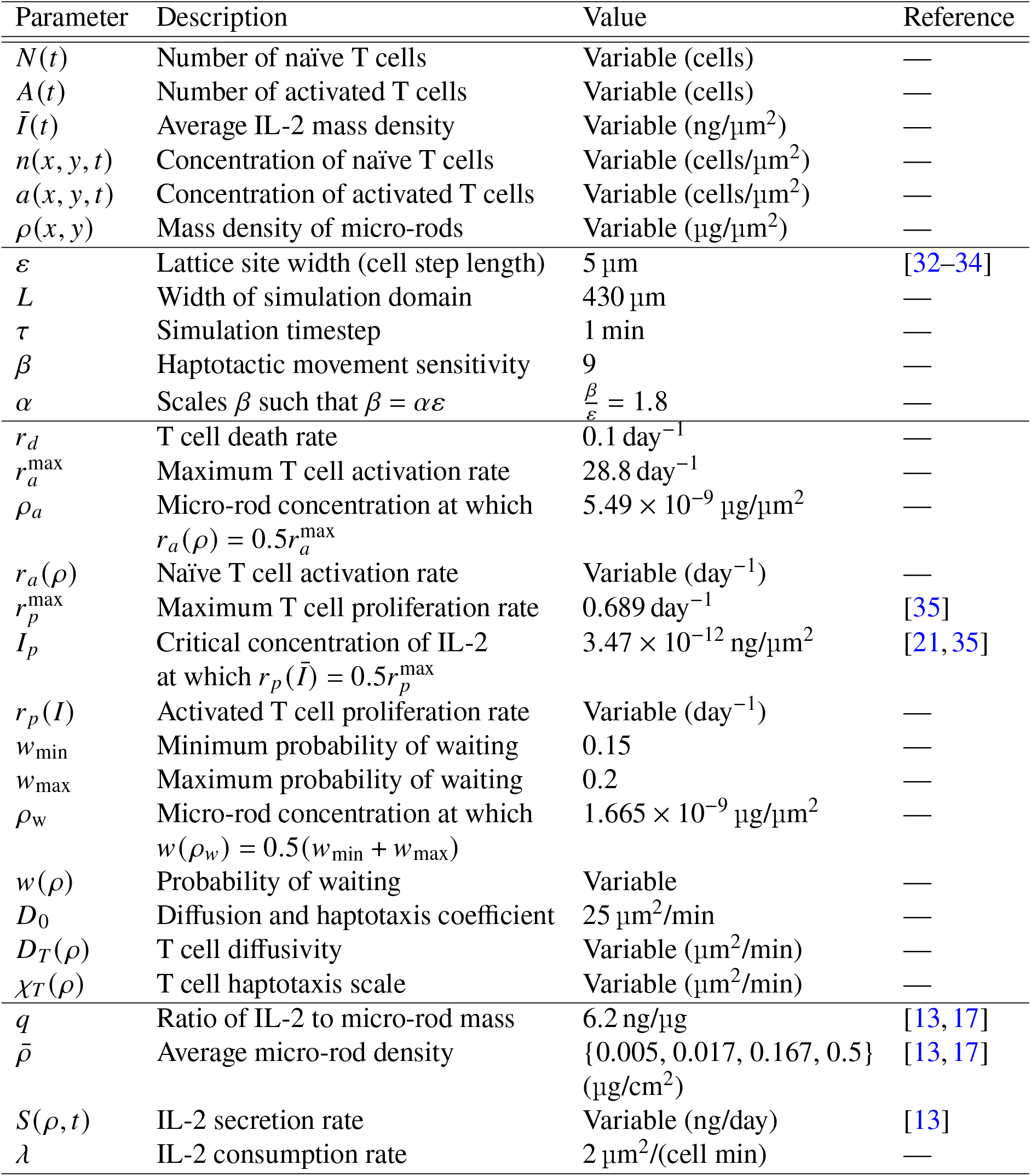
Table of parameters.

**Figure 2.**
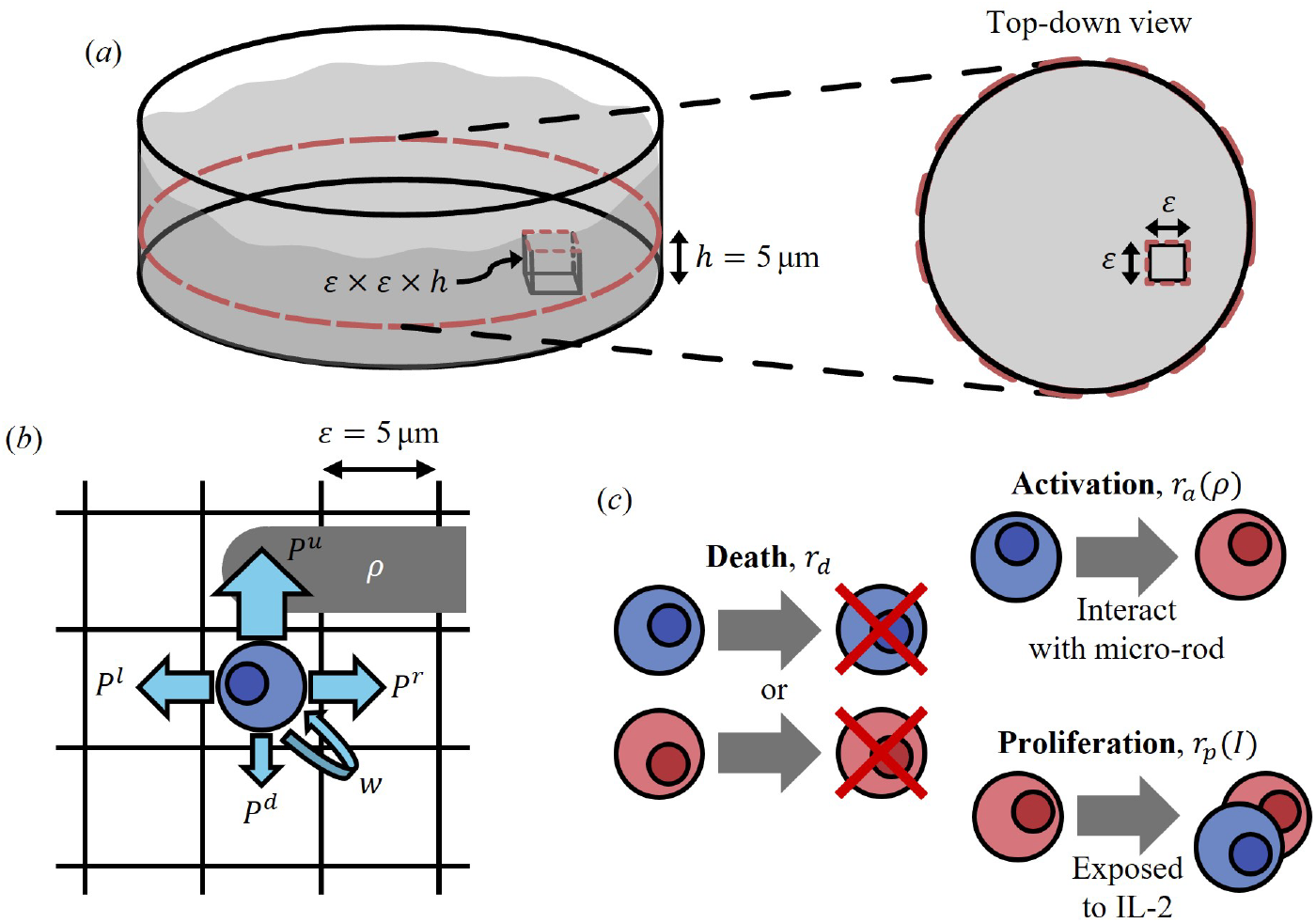
Summary of the computational model design. *(a)* Diagram of the physical interpretation of our 2D simulation domain (right) as a slice out of the culture media (left), with one lattice site shown (not to scale). *(b)* Diagram of T cell movement probabilities on a 2D square lattice with a micro-rod (dark grey). *(c)* Other possible actions for naïve (blue) and activated (red) T cells.

IL-2 is an important factor in T cell expansion, as it promotes the proliferation of activated T cells. In the experiments of Cheung *et al*., micro-rods are fabricated such that IL-2 secretes from within pores in the rods [13] (Fig. 1c). IL-2 molecules are much smaller and densely populated than T cells, so we consider a continuous, time-dependent concentration *I* (*x, y, t*) (ng/µm^2^), governed by the following reaction-diffusion partial differential equation:

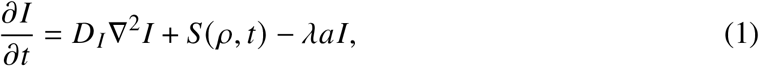

where *D*_*I*_ is the diffusion constant of IL-2 (µm^2^/min), *S* (*ρ, t*) is the total concentration of IL-2 introduced via secretion from micro-rods per unit time (ng/(µm^2^ min)), and *λ* is a rate parameter of IL-2 consumption (µm^2^/(cell min)) by activated T cells which have local concentration *a* (*x, y, t*) (cells/µm^2^). Here we assume that IL-2 in this model does not decay by means other than consumption by activated T cells.

The diffusivity of IL-2 in these micro-rod scaffolds is relatively large, estimated to be approximately 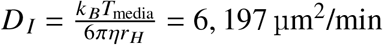 using the Stokes–Einstein–Sutherland equation, which relates the diffusivity and size of IL-2 to the temperature and viscosity of the culture media [36]. We assume that the average micro-rod density is low enough that the presence of micro-rods does not affect the Stokes–Einstein–Sutherland model. Here, *k*_*B*_ = 1.380649 × 10^−23^ m^2^ kg s^−2^ K^−1^ is the Boltzmann constant, *T*_media_ = 20 ^°^C is the temperature of the culture media [17], *η* = 0.958 mPa s is the dynamic viscosity of the culture media [17, 37], and *r*_*H*_ = 2.17 nm is the hydrodynamic radius of IL-2 (converted from the molecular mass of IL-2 of 15.5 kDa [17]). Over our computational domain of area *L*^2^ = 184, 900 µm^2^, the characteristic diffusion time is 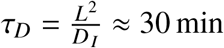, which is insignificant compared to the typical cell culture duration of around two weeks. Therefore, we can assume that the distribution of IL-2 is homogeneous throughout the simulation despite the heterogeneity in its secretion and consumption. This means that we do not consider the spatial distribution of micro-rods or activated T cells to simulate the secretion or consumption of IL-2. Instead of Eq. (1), we simulate the evolution of the average concentration of IL-2, *Ī*(*t*), using a mean-field approximation of the PDE in Eq. (1):

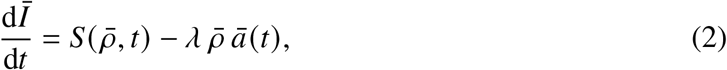

where

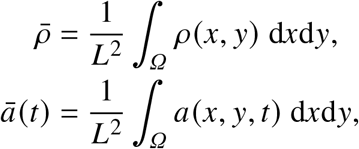

are the spatial averages of micro-rods and activated T cells in our domain, *Ω* : [0, *L*] ^2^, at time *t*, respectively. See Sect. 2.3 for more details on the derivation of a similar ODE. In Sect. 2.4, we use experimental data to define and calibrate the non-spatial secretion rate, 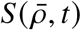.

Even though micro-rods and IL-2 are defined deterministically in our model, T cells are considered to behave stochastically as individuals. T cells in our hybrid agent-based model may be either naïve or activated, and their behaviour changes depending on this state. Each lattice site may contain any amount of naïve T cells and activated T cells. We assume that effector and helper T cells exhibit the same behaviours in micro-rod scaffolds, so we do not differentiate between these cell types. Time is discretised such that lattice site occupancies are updated during timesteps with duration *τ* (in min). In this case, our model predicts T cell positions at discrete times, *t*_*k*_ = *k τ*, where *k* ∈ ℕ. The stochastic variables that govern the number of naïve and activated T cells at some lattice site (*i, j*) and time *t*_*k*_ is 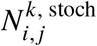 and 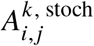, respectively. During each timestep and at each site, the concentration of IL-2 is updated by simulating its evolution using a numerical discretisation of Eq. (2) (see Sect. 2.5 for more details), and the number of T cells is updated by simulating their random motion and actions performed by individual T cells at the previous discrete time. We assume that T cells move according to Brownian motion and haptotaxis along micro-rods, which is the tendency for cells to move towards (and remain near) substrate-bound stimuli [31, 38, 39] (see Fig. 2b). T cell migration is estimated to be on average anywhere between 5 to 15 µm/min *in vivo* [32–34], however, we assume that T cells move more slowly *ex vivo* in these scaffolds due to obstructions by micro-rods. We assume that T cells move by at most one lattice site in each timestep, so a timestep duration of *τ* = 1 min is chosen such that T cells move at an average speed of around 5 µm/min.

We assume that, in the experiments of Cheung *et al*., T cells are attracted towards the *α*CD28 and pMHC molecules bound to lipids on the micro-rods via haptotactic motion. In our model, T cells move preferentially towards regions of high micro-rod density, similarly to existing models of cell chemotaxis [27, 30]. T cells may also become quiescent and wait in place due to the presence of micro-rods, with probability *w* (*ρ*_*i, j*_). Given that a cell at a lattice site (*i, j*) and time *t*_*k*_ is selected to undergo motion with probability *M*, the probabilities for moving left to (*i* − 1, *j*), right to (*i* + 1, *j*), downwards to (*i, j* − 1), and upwards to (*i, j* + 1) respectively are

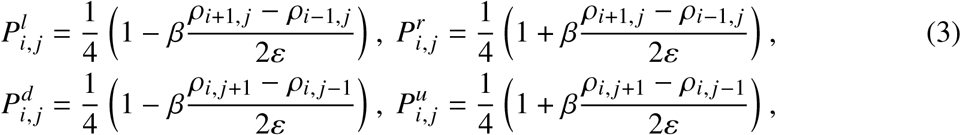

where *β* scales the strength of haptotaxis and needs to be selected small enough so that each movement probability is positive and no greater than 1. Central differences of *ρ* are used to bias movement in the direction of increasing micro-rod density (Fig. 2b), and to ensure that these movement probabilities sum to 1. We also see that 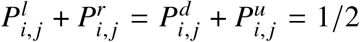, which asserts that T cells have equal probability of moving left or right as they do of moving up or down. Intercellular interactions such as cell-to-cell adhesion and volume filling effects are not included as it is assumed that they do not impact T cell expansion significantly in these micro-rod scaffolds. Additionally, multiple T cells in one lattice site may represent cells that are offset vertically, so we assume that these cells do not meaningfully interact.

If T cells do not move or they become quiescent, they may perform other actions depending on their activation state (Fig. 2c). Naïve T cells may activate when they interact with the stimuli attached to micro-rods, which is modelled as a rate of activation depending on the local micro-rod density, *r*_*a*_ (*ρ*_*i, j*_) (probability per unit time). We assume that T cells do not exhibit a range of intermediate activation states, so that a naïve T cell becomes fully activated when it undergoes activation. Activated T cells may proliferate if they are exposed to a sufficient concentration of local IL-2, which is modelled as an IL-2-dependent proliferation rate *r*_*p*_ (*Ī*) (probability per unit time). Here we assume that proliferation is asymmetric and produces a new naïve daughter cell at the same location as the proliferating activated cell. In either state, T cells are assumed to die at a constant rate, *r*_*d*_. We assume that the dependence of these rates upon micro-rod-bound stimuli and *Ī* is described by Michaelis-Menten reaction kinetics [40], such that *r*_*a*_ (*ρ*) and *r*_*p*_ (*Ī*) are given by

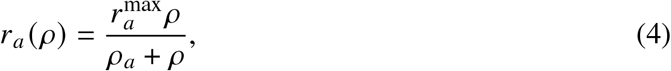

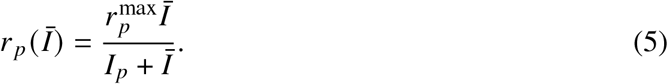

In Eqs (4) and (5), 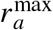 and 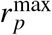 describe the limiting rates for activation and proliferation, and *ρ*_*a*_ and *I*_*p*_ correspond to the concentrations of *ρ* and *Ī* at which *r*_*a*_ and *r*_*p*_ are half their maximal values, respectively. The parameters involved with *r*_*p*_ (*Ī*) were fitted to data (see Sect. 2.4). Similarly, we assume that the probability of waiting, *w*(*ρ*), follows Michaelis-Menten kinetics:

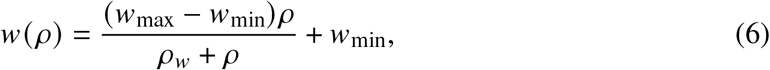

where *w*_min_ is the minimum probability of waiting (in the absence of micro-rods), *w*_max_ is the maximum probability of waiting, and *ρ*_*w*_ is the micro-rod concentration at which the probability of waiting is halfway between its minimum and maximum values. We select *ρ*_*w*_ to be the average local micro-rod concentration in a 333 µg/mL scaffold, which is a concentration commonly used in experiments [13, 17].

See Sect. 2.5 for details on the numerical simulation of the ABM. During each time increment, *τ*, each cell is selected to undergo either death with constant probability *τr*_*d*_, activation with probability *τr*_*a*_ (*ρ*_*i, j*_) if the cell is naïve, or proliferation with probability *τr*_*a*_ (*Ī*) if the cell is activated. Since we assume that cells may only move, wait in place, or perform one action in one timestep, the probability of moving for naïve cells is *M* (*ρ*_*i, j*_) = 1 − *w*(*ρ*_*i, j*_) − *τr*_*d*_ − *τr*_*a*_ (*ρ*_*i, j*_), the probability of moving for activated cells is *M* (*ρ*_*i, j*_, *Ī*) = 1 − *w*(*ρ*_*i, j*_) − *τr*_*d*_ − *τr*_*p*_ (*Ī*). We assume that *τ* or the limiting reaction rates for *r*_*d*_, *r*_*a*_ (*ρ*) and *r*_*p*_ (*Ī*)are small enough that *M* > 0 in both cases. The local concentration of naïve and activated T cells at some lattice site is given by the local number of cells divided by the area of the site, 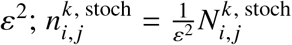 and 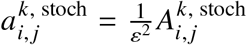 respectively. Our simulation domain represents a small *L* × *L* portion of the whole culture dish and we assume that the rest of the culture dish has a similar distribution of micro-rods and T cells, so we apply periodic boundary conditions on T cell movement.

### 2.2 Continuum limit

The continuum limit of the behaviour of T cells in the ABM provides us with a deterministic and continuous model of the average T cell and IL-2 concentrations over time, which can be useful for computational efficiency and analysis. A continuum limit can be derived by considering that the discrete spatial step (in this case the T cell movement step size, *ε*) and timestep, *τ*, both approach zero [41–43]. We first describe the expected or average evolution of the number of naïve cells at lattice site (*i, j*) and time *t*_*k*_ in a deterministic manner by considering 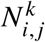 as the expected value of the stochastic variable 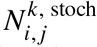 [44]. The following equation describes the balance of naïve cell numbers at an interior site (*i, j*) due to naïve cell movement, death, activation and production by asymmetric cell division from activated T cells:

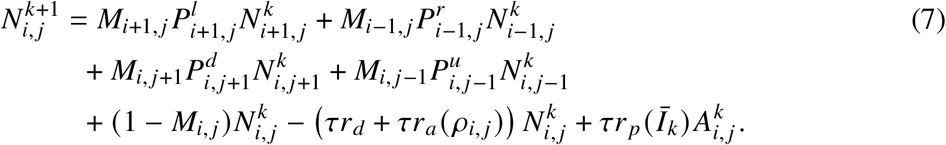

Here, *M*_*i, j*_ = *M* (*ρ*_*i, j*_) is the probability of moving, and 1 − *M*_*i, j*_ is the probability of remaining stationary, either due to quiescence or preoccupation while performing other actions. We assume that this expected evolution for naïve cells in site (*i, j*) occurs simultaneously with the evolution for activated cells and IL-2, such that, for example, activated cell death or movement at time *t*_*k*_ can only impact the production of naïve cells in the next time, *t*_*k*+1_. Eq. (7) may also be used to describe the average discrete evolution of naïve cell concentration, 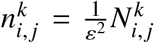, and similarly for activated cell concentration, 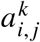. Dividing Eq. (7) by the area of the lattice site (*i, j*), substituting the equations for movement probabilities (Eqs (3)) and rearranging, we have

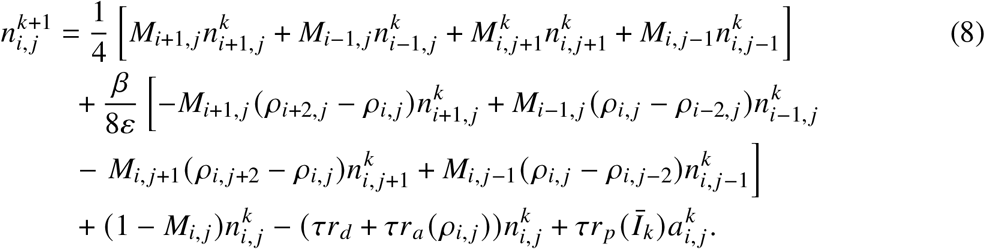

We now introduce continuous functions in time and space, *n* (*x, y, t*), *a* (*x, y, t*) and *ρ* (*x, y*), which describe the average concentrations of naïve T cells, activated T cells and micro-rods in the continuum limit, respectively, such that 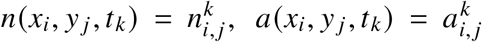, and *ρ*(*x*_*i*_, *y*_*j*_) = *ρ*_*i, j*_. Replacing the discrete concentrations of naïve and activated T cells, micro-rods and IL-2 with their continuous definitions, we Taylor expand all terms in the above equation about (*x*_*i*_, *y*_*j*_, *t*_*k*_). For the term on the left-hand side of Eq. (8), we see that

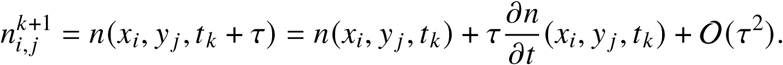

After Taylor expanding, all terms are evaluated at *x*_*i*_, *y*_*j*_, *t*_*k*_, so we drop this notation for conciseness. Similarly, for terms on the first line of the right-hand side of Eq. (8),

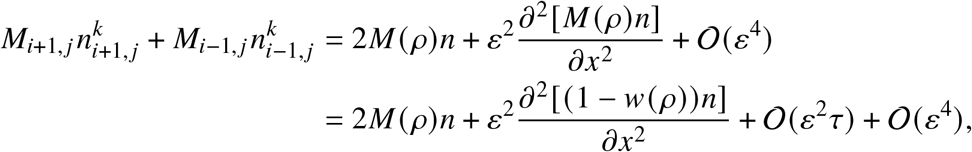

where the second equality uses the fact that

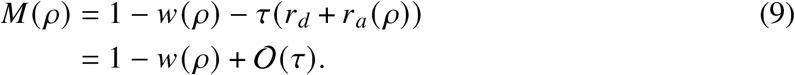

By similar reasoning, terms on the second and third lines of the right-hand side of Eq. (8) are expanded as

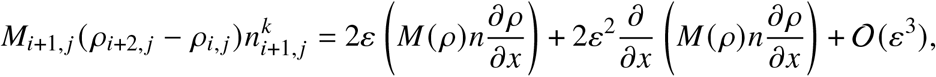

so that

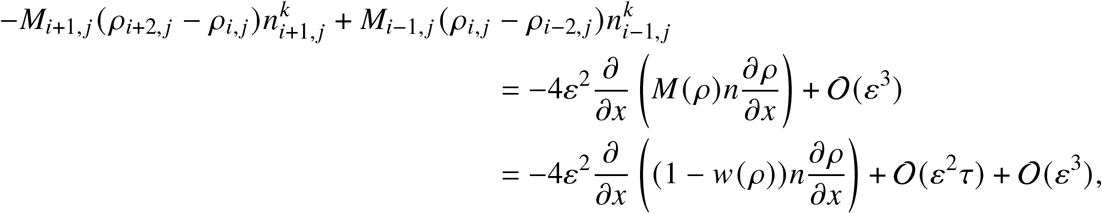

where the last equality again applies Eq. (9). Using these expansions along with similar Taylor expansions for the remaining terms, Eq. (8) becomes

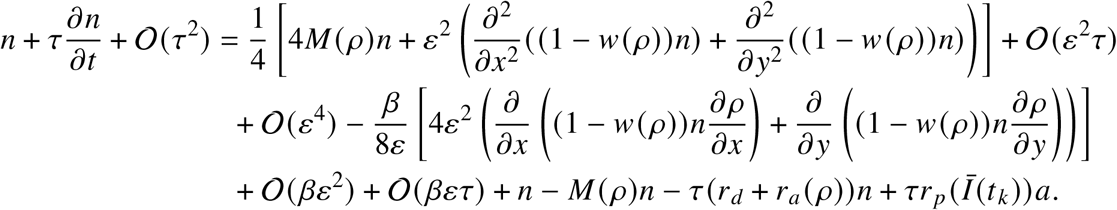

Cancelling terms, dividing through by *τ*, and rearranging, gives

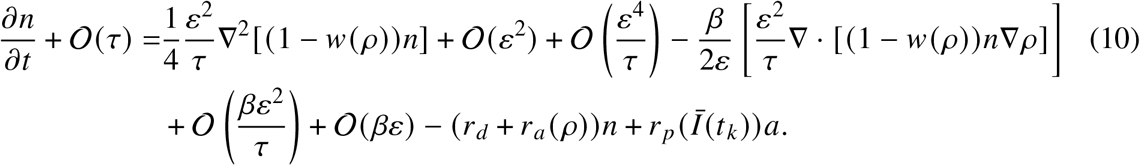

We consider the limit as *ε, τ* → 0, assuming that 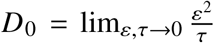 exists and is finite, and also that *β* = *αε* (the strength of T cell haptotaxis is proportional to their step length, *ε*) [41]. Under these assumptions, as 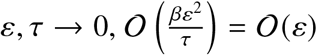 and 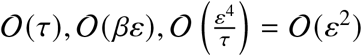 are all subdominant. Taking this limit of Eq. (10), we obtain

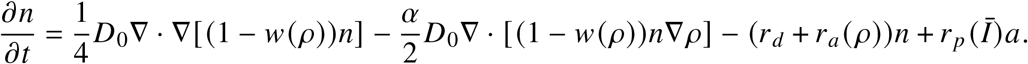

Finally, using 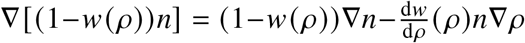, the above equation can be rearranged to obtain a PDE which describes the expected evolution of the concentration of naïve T cells:

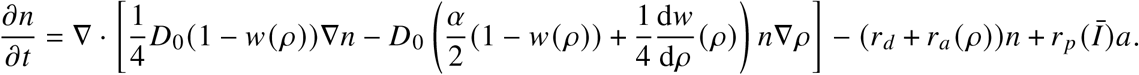

This process is repeated to formulate a PDE that governs the concentration of activated T cells, which exhibits the same diffusion and advection, but differs in its reaction terms (activated cell concentration increases due to activation of naïve cells and reduces due to cell death). The ODE governing IL-2 in Eq. (2) depends on the spatial average of activated cells, ā (*t*), which is the integral of the mass concentration of activated cells, *a*, over the simulation domain, *Ω* : [0, *L*] ^2^, divided by the area of the domain. The following is the complete continuum limit model, which consists of a nonlinear system of differential equations involving PDEs which govern naïve and activated T cells, and an ODE which governs the average concentration of IL-2:

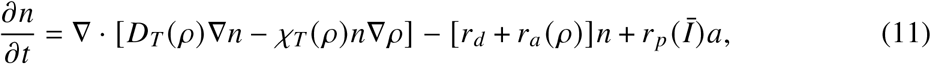

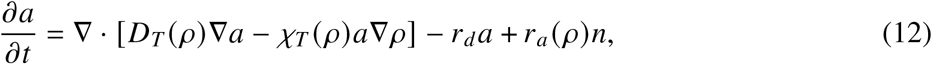

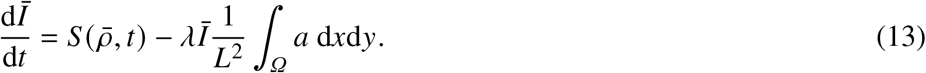

In Eqs (11) and (12),

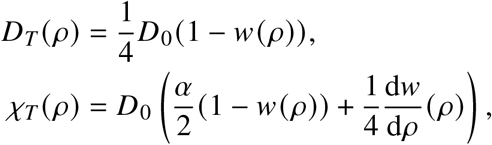

represent T cell diffusion and the haptotactic scale (scales the strength of cell motion in the direction of ∇*ρ*) respectively. This model is solved numerically subject to periodic boundary conditions, as detailed in Sect. 2.5. Note that, for small values of *w* (*ρ*) or large values of *α* or 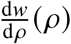, this model may experience finite time blow-up similar to computational models which describe cell chemotaxis [27, 30].

### 2.3 Mean-field approximation

In order to model the evolution of T cell populations directly, we consider an analogous non-spatial model derived from the PDE model by applying a mean-field approximation. A similar procedure was used in Sect. 2.1 to generate the ODE governing the average concentration of IL-2 in Eq. (2). Consider the total numbers of naïve and activated T cells in the domain, *N* (*t*) and *A*(*t*) respectively, defined by

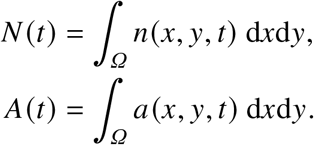

Integrating the PDE governing naïve T cells (Eq. (11)) over space gives

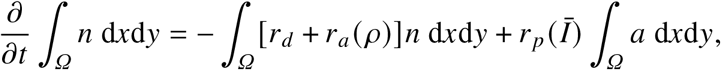

since the transport terms give no contribution by application of the divergence theorem with periodic boundary conditions. The mean-field approximation assumes that micro-rods are uniformly distributed throughout the domain, such that *ρ* can be replaced by its spatial average, 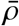:

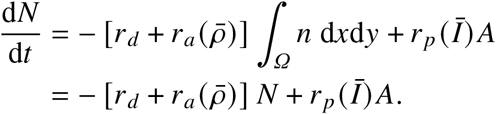

Repeating this procedure for activated T cells and including the ODE for IL-2, we obtain the nonlinear system of ODEs:

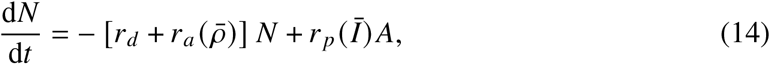

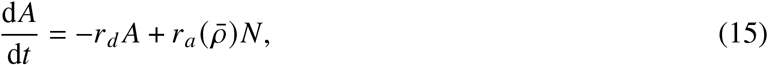

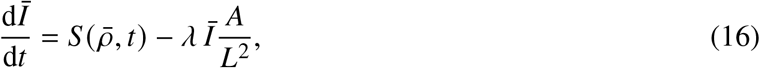

where Eq. (16) is equivalent to Eqs (2) and (13). We solve this ODE system numerically (see Sect. 2.5).

### 2.4 Model fitting to experimental data

The IL-2-dependent proliferation rate, *r*_*p*_ (*Ī*) in Eq. (5), has parameters for the maximum proliferation rate, 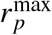, and the IL-2 concentration at which the proliferation rate is half the maximum, *I*_*p*_. We fit these parameters to data describing the total expansion of T cells under constant concentrations of IL-2. Fig. 3a shows the expansion of T cells under a constant IL-2 concentration of 1.11 ng/mL from experiments by Kaartinen *et al*. in 2017 [35], with data digitised using Web-PlotDigitiser [45]. Here we assume that T cells do not die and that all T cells in this experiment are always activated such that they may always proliferate, and that IL-2 is not consumed during expansion. Under these assumptions, the total number of cells, *N*_*T*_ (*t*), is governed by the ODE: 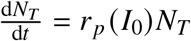, where *r*_*p*_ (*I*_0_) is the T cell proliferation rate under a constant concentration of IL-2, *I*_0_. This implies that 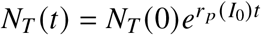, or the fold expansion 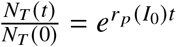.

**Figure 3.**
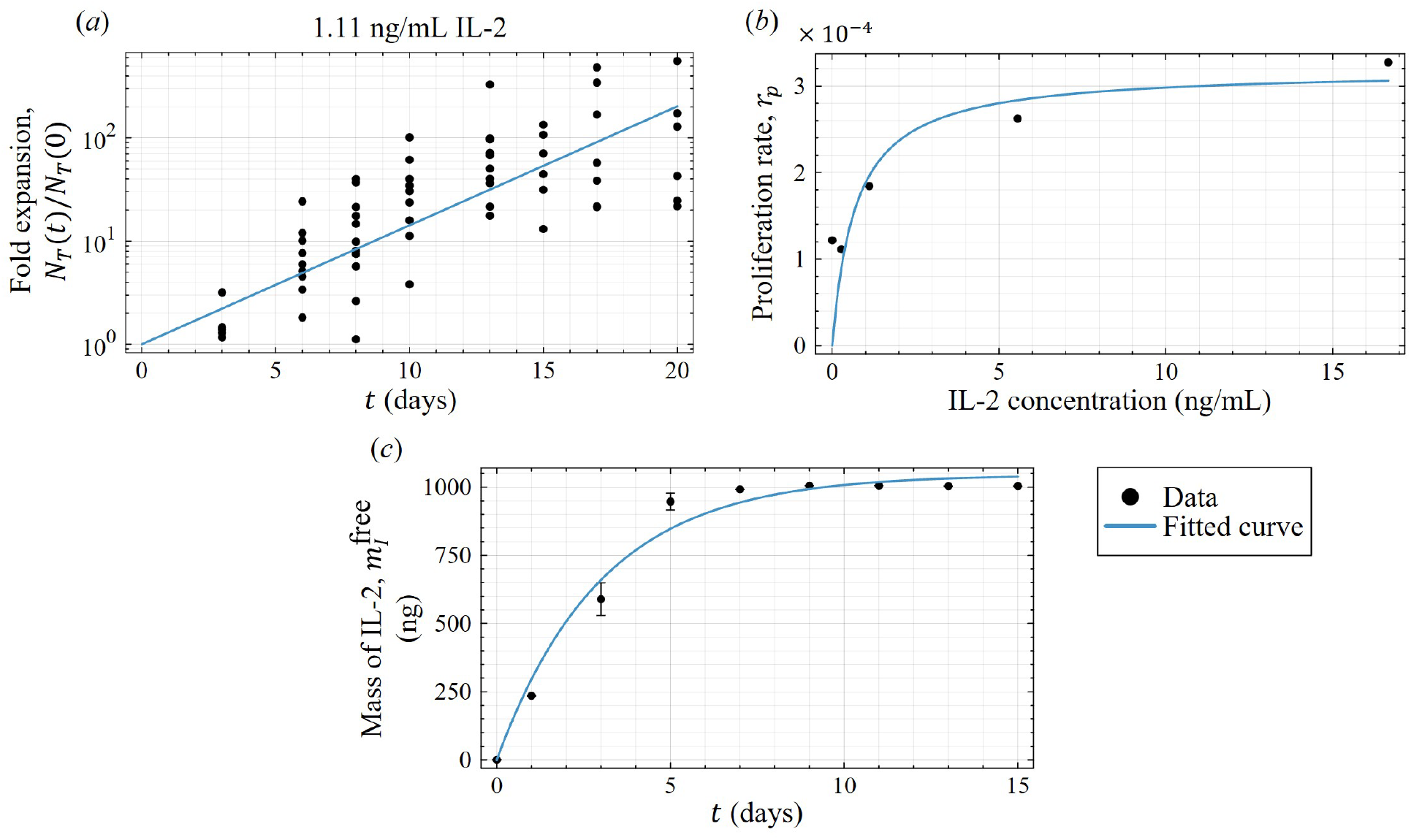
Model fitting for *(a,b)* activated cell proliferation rate and *(c)* IL-2 secretion from micro-rods. *(a)* Exponential fit of T cell expansion used to recover proliferation rate when IL-2 concentration is 1.1 ng/mL (data reproduced from [35]). *(b)* Michaelis-Menten equation fitted to the proliferation rates at varying IL-2 concentrations (recovered from analysis in *(a)* and similar in Appendix A). *(c)* Exponential fit of the total mass of free IL-2 in the system in the absence of T cells (data reproduced from [13]).

We fit *r*_*p*_ (*I*_0_) to five data sets detailing T cell expansion under various constant concentrations *I*_0_ of IL-2 (Fig. 3a and Fig. 9 in Appendix A). The data points in Fig. 3b represent these fitted proliferation rates. We then fit the two parameters of Eq. (5), 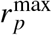 and *I*_*p*_, such that *r*_*p*_ (*Ī*) approximately passes through these proliferation rates under the constant IL-2 concentrations *I*_0_ (Fig. 3b). We find that 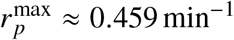 and *I*_*p*_ ≈ 3.47 × 10^−12^ ng/µm^2^. Our fitted model suggests that the T cell proliferation rate increases quickly when the concentration of IL-2 is below or around *I*_*p*_, and approaches the maximum rate after *I*_*p*_, which is roughly supported by experiments in [21]. Since our simulation domain represents a *h* = 5 µm tall horizontal slice of a culture dish, we convert mL to the volume occupied above some area in our domain. Specifically, *h* × 10^−12^ mL corresponds to the volume above 1 µm^2^ of surface area in our 2D model. Since these parameter values were fitted under the assumption that T cells do not die in these experiments, but cell death was likely occurring to some degree, the true proliferation rate can be expected to be larger than that fitted to the data. We assume the maximum proliferation rate in our model is 150% of the maximum rate fitted to this data, so that 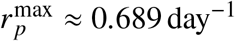.

We also fit the total secretion rate of IL-2 from micro-rods, 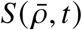, using data (digitised from [13] with WebPlotDigitiser [45]) which describes the total amount of IL-2 released by micro-rods over time in the absence of T cells, such that IL-2 is not consumed (see Fig. 3c) [13]. We first assume that IL-2 molecules secrete from a single micro-rod with constant probability per unit time, *v*, such that the total mass of IL-2 stored inside micro-rods in the domain, 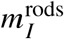, and the total mass of free IL-2 (not inside micro-rods), 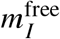, are governed by the ODEs:

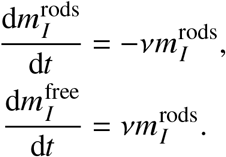

Assuming that there is no free IL-2 initially, we see that 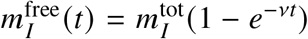, where 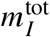 is the total initial mass of IL-2 loaded inside all the micro-rods. Fitting both *v* and 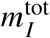 to data (Fig. 3c) gives *v* ≈ 0.33 day^−1^ and 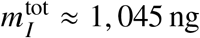. The average concentration of free IL-2 in the domain is 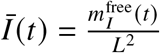, so comparing the ODE governing 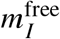 to Eq. (2) in the absence of IL-2 consumption by T cells,

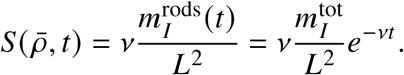

This equation can be reformulated to describe the secretion of IL-2 in terms of the ratio of loaded IL-2 mass to micro-rod mass, 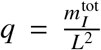, and the average micro-rod concentration, 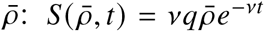. In practice, micro-rods are commonly prepared such that around 3,100 ng of IL-2 is loaded into (and retained inside) 500 µg of micro-rods [13, 17]. As such, in our model, we use the ratio of IL-2 mass to micro-rod mass, *q* = 6.2 ng/µg. Also, scaffolds with an average micro-rod concentration of 333 µg/mL are commonly used [13, 17], so we select 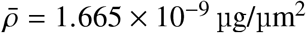, unless otherwise stated.

Parameters *r*_*d*_, *λ* and *α*, and those involved in the functions *r*_*a*_ (*ρ*) (Eq. (4)) and *w* (*ρ*) (Eq. (6)), are chosen arbitrarily due to a lack of data or information on the behaviours of T cells in micro-rod scaffolds. See Table 1 for these parameter values.

### 2.5 Numerical simulation

All simulations and analyses are performed in Julia (v1.11.2).

We simulate micro-rod scaffolds by randomly generating the micro-rod mass concentration for each lattice site, *ρ*_*i, j*_. To determine the number of micro-rods in our domain, we consider the average concentration of micro-rods and the volume represented by our lattice. Our simulation domain has surface area *L*^2^ = 184, 900 µm^2^ and it represents volume with height *h* = 5 µm, so it is analogous to a culture media volume of *L* × *L* × *h* = 0.925 nL. Given the average concentration of micro-rods, 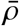, the total mass of micro-rods in the domain is 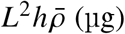, which is divided by the mass of an individual micro-rod to determine the number of rods. For each micro-rod, we randomly generate its length, *l*, and diameter, *d*, assuming that they are uniformly distributed on the intervals [38.7, 102.1] (µm) and [3.9, 6.3] (µm), respectively, which are the dimensions of experimentally fabricated micro-rods [13]. We then randomly sample the coordinates of the centre of the micro-rod, *x*_*c*_, *y*_*c*_, and its rotation angle, *θ*, from uniform distributions (*X, Y*) ~ (*U* (0, *L*), *U* (0, *L*) and *Θ ~ U* (0, *π*), respectively, unless otherwise stated. We consider the line that passes through (*x*_*c*_, *y*_*c*_) with rotation angle *θ* measured anti-clockwise from the positive *x* axis. For all lattice sites whose centres are within *d* 2 (µm) of this line, if the distance from the centre of the lattice site to (*x*_*c*_, *y*_*c*_) is less than or equal to *l* / 2 (µm), we increment the micro-rod density at this lattice site by the mass of one micro-rod divided by *ld* (µm^2^). We also apply periodic boundary conditions such that micro-rods which extend beyond the simulation domain are mapped to the opposite side of the domain.

T cells and IL-2 are simulated by iterating through all timesteps (with a duration of *τ* = 1 min) until 14 days (or longer), as this is the typical duration of experiments involving T cell expansion in micro-rod scaffolds [13,17]. Initially, there are no activated T cells and no free IL-2; all the IL-2 is loaded inside micro-rods. We randomly place 10 naïve T cells in the *L* × *L* simulation domain according to a uniform distribution. Assuming that there is a similar average concentration of naïve T cells in all other slices of the culture media, then the whole culture media would be seeded with around one million naïve T cells, which is similar to experiments [13, 17]. During each timestep and for each T cell (selected sequentially at random without replacement), a random number, *r U ~* (0, 1), is sampled and compared to the probabilities of performing each individual action. Only one action is performed per timestep. Probabilities involving *ρ* or *Ī* are evaluated within the lattice site of the current cell, (*i, j*), at the time of the previous iteration, *t*_*k*_.

We first determine whether the current T cell dies in this timestep by checking if *r* < *τr*_*d*_. If so, then that cell is removed from the simulation at time *t*_*k*+ 1_, otherwise, if the cell does not die, then we continue checking the other possible actions. If the T cell is naïve and *τr*_*d*_ ≤ *r* < *τ*(*r*_*d*_ + *r*_*a*_ (*ρ*_*i, j*_)), then it becomes an activated T cell at time *t*_*k* + 1_. Otherwise, if the T cell is activated, we perform similar comparisons with the probability of proliferating, *τr*_*p*_(*I*_*k*_). If proliferation occurs, a new naïve T cell is introduced at lattice site (*i, j*) and time *t*_*k*+ 1_. If the current T cell does not perform any of these actions, then we check if it remains quiescent with probability *w* (*ρ*_*i, j*_), and if so, it does not change state or move until the next timestep. Otherwise, movement probabilities in Eqs (3) are compared to *r*, and the T cell moves in the selected direction such that it occupies that neighbouring site at time *t*_*k*_. We apply periodic boundary conditions as we assume that other regions of the culture dish have a similar composition of micro-rods and T cells. The sequence of checks required to simulate T cell movement and reactions is not important and may be performed in any order, since cells are only able to perform one action in each timestep. The average concentration of IL-2 at the next timestep, *Ī*_*k* + 1_, is found by numerically iterating the ODE in Eq. (2) with the fourth order Runge-Kutta method using the DifferentialEquations.jl package [46]. This procedure is then repeated in the next timestep at time *t*_*k*+ 1_.

To determine the average behaviour of T cells or the average concentration of IL-2 over time, this simulation is repeated multiple times and the results can be collated. To compare our results with the experimental results obtained by Cheung *et al*., we scale T cell populations and IL-2 mass to represent the expansion that we expect to see in the whole culture dish. Zhang *et al*. describe that they initially expand T cells using 100 nL of media containing micro-rods [17], so we assume our simulation domain represents a small proportion of this media. Since our simulation domain represents a volume of 0.925 nL, we scale our results by multiplying the number of T cells and mass of IL-2 by 100/0.925.

The PDEs governing T cells (Eqs (11) and (12)) are discretised in space using the finite volume method (FVM) with periodic boundary conditions and the same square lattice as used in the ABM, and discretised in time using the Crank-Nicolson method with timestep duration *τ* = 1 min. The concentration of naïve T cells is initially uniform and corresponding to a total of 10 cells, and the concentrations of activated T cells and IL-2 are initially zero. These PDEs and the ODE governing IL-2 (Eq. (13)) are solved to find the concentrations of naïve T cells, activated T cells and IL-2 at each discrete time. The IL-2 ODE involves the spatial average of the concentration of activated T cells, which is numerically estimated by averaging the concentrations at each lattice site at time *t*_*k*_. See Appendix B for more details regarding the numerical methods used to solve the PDEs governing T cells.

## 3 Results

### 3.1 Stochastic simulations and comparison to experiments

We start by showing numerical results of an ABM simulation representing a typical experiment using a 333 µg/mL micro-rod scaffold (Fig. 4). To match the initial concentration of T cells used in experiments, we randomly distribute 10 naïve T cells into our 430 µm ×430 µm simulation domain according to a uniform distribution (Fig. 4a). The initial fast release of IL-2 from micro-rods means that the mass of IL-2 in the culture media first increases (Fig. 4c). This allows cells which have been activated by micro-rods to proliferate, causing rapid expansion in the activated T cell population (Fig. 4b). As a result, IL-2 is quickly consumed by activated cells, and the total mass of IL-2 in the domain decreases after around 1 day. Eventually, there is insufficient IL-2 to support rapid proliferation, which causes the naïve T cell population to peak at around 5 days. A few days later, the activated T cell population also reaches a maximum as there is not enough activation of naïve T cells.

**Figure 4.**
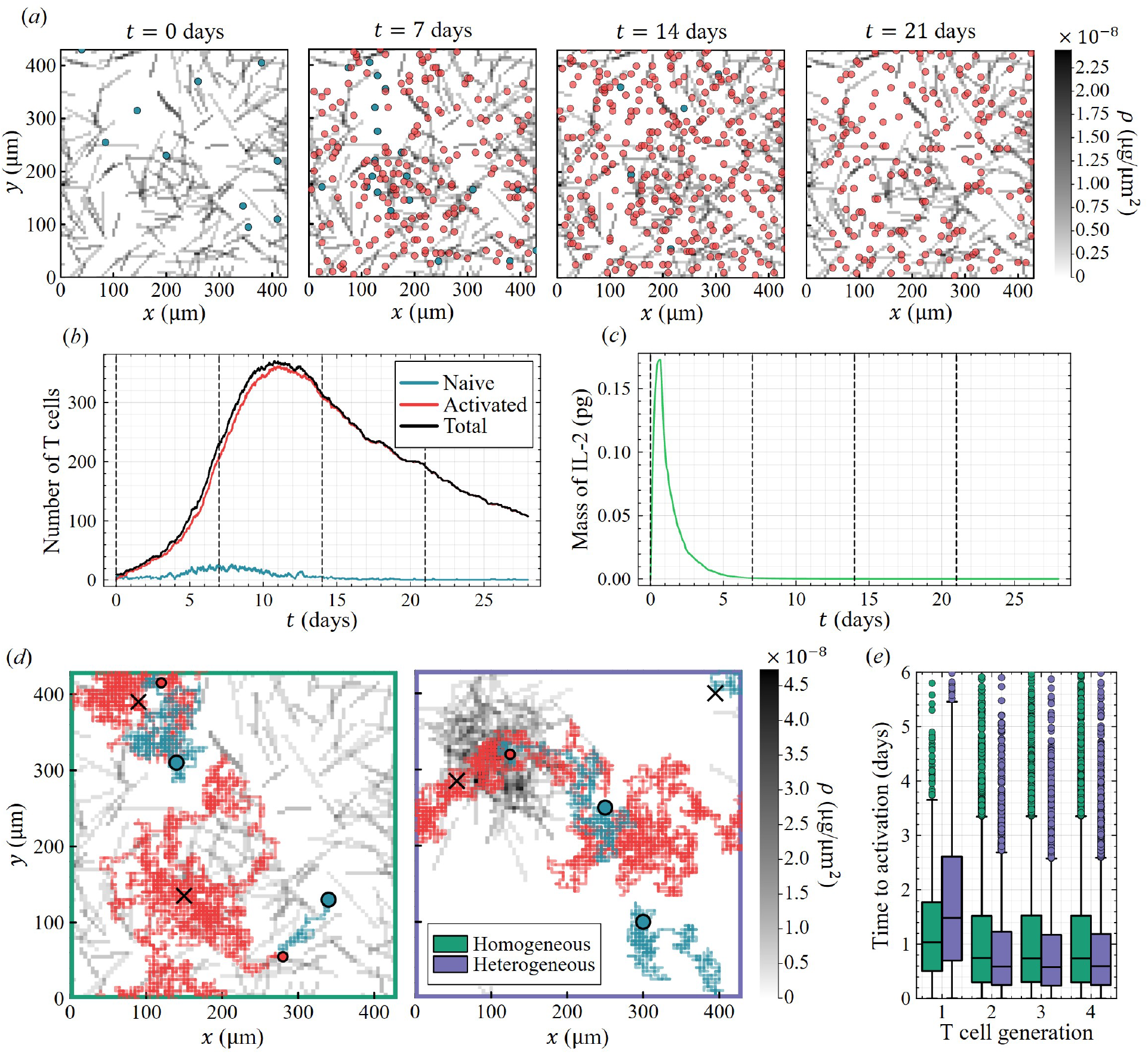
Representative stochastic realisation using a randomly generated 333 µg/mL microrod scaffold showing *(a)* naïve (blue) and activated (red) T cells overlayed on a heatmap of micro-rod concentration (grey) at 0, 7, 14, and 21 days, with *(b)* total populations of T cells, and *(c)* total mass of IL-2 in the simulation domain. *(d)* Representative cell trajectories in a homogeneous scaffold (left) and a heterogeneous scaffold (right), where the cells initially start at the blue circles, become activated at the red circles, and die at the black crosses. *(e)* Median and interquartile range of time taken to activate for each cell generation in the homogeneous and heterogeneous scaffolds shown in *(d)* (data from 50 simulations on each scaffold, omitting some outliers).

Since T cells in our model preferentially move along micro-rods by haptotaxis, and are quickly activated by the stimuli attached to micro-rods, activated T cells outnumber naïve T cells and tend to cluster around regions of high average micro-rod concentration (Fig. 4a). This behaviour is seen experimentally (see Fig. 5a) as T cells collect around micro-rods and eventually form large cell-rod clusters, which reportedly promote rapid expansion [13]. Our model suggests that expansion is driven by activated cells in cell-rod clusters producing new naïve cells that are immediately activated by the micro-rods in the cluster. In the model, increasing the strength of T cell haptotaxis and the probability of cell quiescence around micro-rods contributes to the clustering of T cells around micro-rods, which improves the overall expansion (data not shown). Such changes in haptotaxis might be achieved experimentally by increasing the amount of activating stimuli attached to the surface of micro-rods, which we also expect to increase the T cell activation rate.

**Figure 5.**
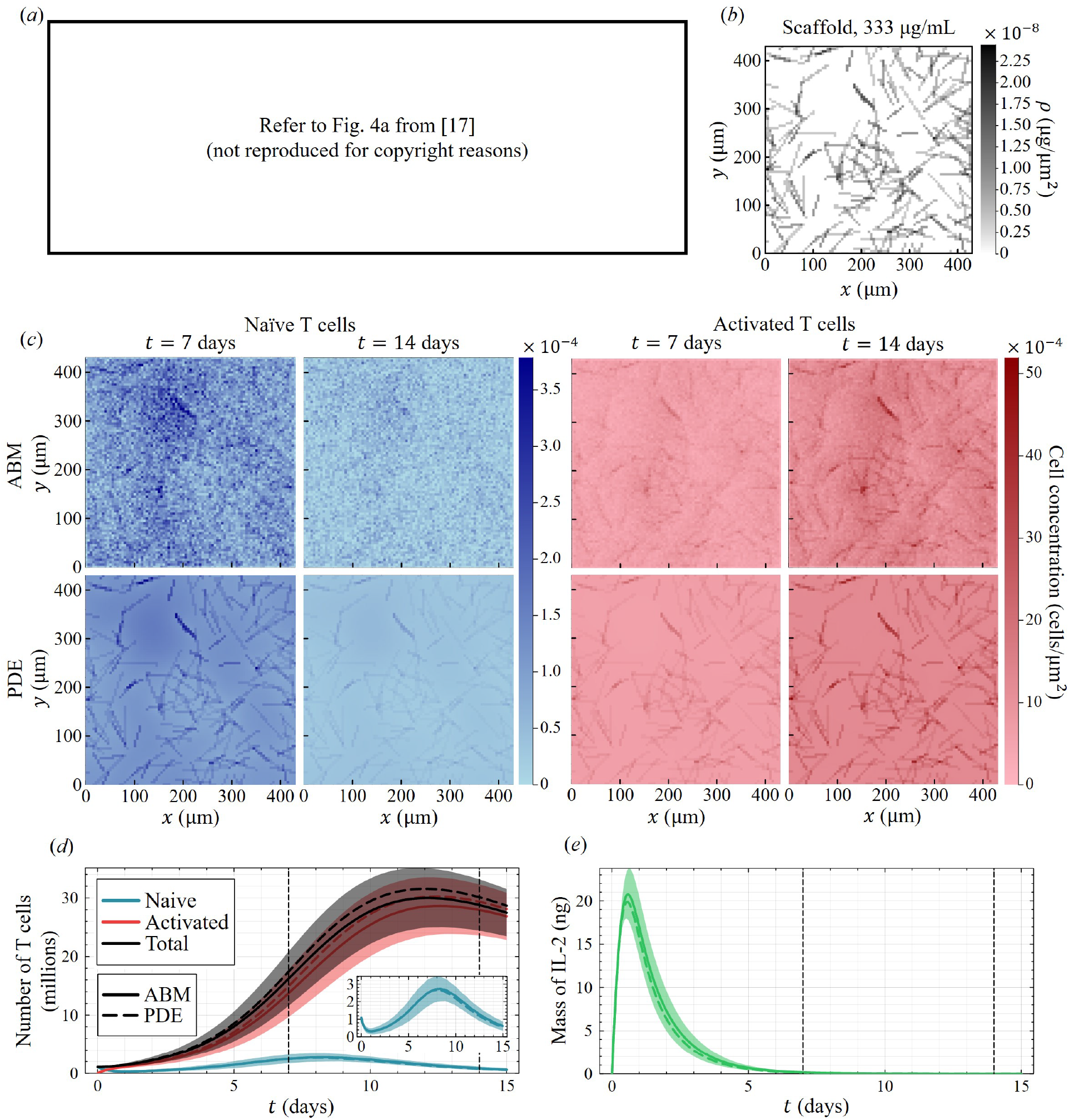
Experimental data and computational simulations from spatial models of T cell expansion with micro-rods. *(a)* Bright-field micrographs of a scaffold before co-culture with T cells (left) and after introducing T cells at 2 hours (middle) and 1 day (right) (400 µm scale bar), reproduced from [17]. *(b)* Micro-rod mass concentration of a representative 333 µg/mL micro-rod scaffold. *(c)* Distributions of naïve (blue) and activated (red) T cells from the ABM (average of 2,000 realisations) and its continuum limit at 7 and 14 days. *(d)* Resulting T cell populations (average and standard deviation of the ABM are shown, with a zoomed view of the naïve cell population) and *(e)* total mass of IL-2 (results scaled up to represent those expected in the whole culture dish).

An advantage of our model is to be able to track the trajectories of T cells and their interactions with the micro-rods. Stochastic simulations in Fig. 4d reveal that, in well-mixed regions of the micro-rod scaffold, T cells are able to locate micro-rods and form these clusters easily. However, in regions of a scaffold devoid of micro-rods (like in Fig. 4d), naïve cells may die before activating, which hinders cluster formation and rapid expansion, as seen by the trajectory of the bottom right cell in Fig. 4d. Fig. 4e shows the average time for an individual cell to become activated based on the cell’s generation (the first generation refers to the initial cells, the second refers to cells which divided from those in the first generation, and so on). T cell activation time is non-normally distributed in both scaffolds (using the Shapiro-Wilk test), so we perform Mann-Whitney U tests and determine that there is significant evidence that activation times differ between scaffolds for each cell generation. Over 50 simulations (with 10 first-generation cells each), we see that initial T cells generally take a longer time to become activated in the heterogeneous scaffold, but future generations of cells are able to activate sooner than those in the homogeneous scaffold. We investigate the impacts of scaffold heterogeneity on T cell expansion further in Sect. 3.5. Simulating expansion over 14 days with the homogeneous scaffold in Fig. 4d takes 2.6 seconds on average, and simulation with the heterogeneous scaffold takes 2.2 seconds on average.

### 3.2 Comparison of stochastic and deterministic frameworks

Fig. 5 summarises the average behaviour of T cells and IL-2 in the scaffold in Fig. 5b when the stochastic simulation in Fig. 4 is repeated over 2,000 times. The average spatial distributions of naïve and activated T cells are compared with the PDE model (Eqs (11)–(13)) at 7 and 14 days in Fig. 5c. The population results in Figs 5d,e show the average and standard deviation of stochastic simulations with the PDE model predictions, where the number of T cells and mass of IL-2 is scaled up to represent those expected from the whole culture dish (as detailed in Sect. 2.5). The PDE model accurately captures the average qualitative behaviours of naïve and activated T cells in the ABM, both in spatial distribution and in population growth. Both of these models predict that activated T cells will cluster around micro-rods on average, as expected from stochastic simulations in Fig. 4a and observations from experiments (Fig. 5a). Naïve T cells are found with relatively large concentrations in regions devoid of micro-rods as they become activated quickly by micro-rods.

Similarly to the patterns seen in Fig. 4, we see that the average concentration of IL-2 in Fig. 5e declines to approximately *I*_*p*_ at 8 days, which is the concentration at which the T cell proliferation rate is half its maximum. As the IL-2 concentration continues to decrease past this critical concentration, T cell death begins to dominate proliferation. This causes a decline in the naïve T cell population and a decreased overall growth rate of activated T cells (Fig. 5d). In this scaffold, both models predict that the total population of T cells peaks at around 30-fold expansion (total number of T cells divided by the initial number).

The ABM model allows us to estimate uncertainty in the simulation results. The standard deviations of the populations of T cells and IL-2 (shown as shaded regions in Fig. 5d,e) indicate that there is some variability in the total expansion of T cells over time. Experimental results suggest similar variability relative to the total expansion of T cells in these micro-rod scaffolds [13]. Compared to the ABM, the PDE model slightly overestimates activated T cell expansion, and therefore underestimates the total mass of IL-2 due to increased consumption. The averages of T cell population predictions from the ABM display slower growth compared to the PDE model due to some stochastic realisations in which all T cells became extinct. However, the PDE model is much more computationally efficient (and may be made faster by increasing the simulation timestep), so given that it is sufficiently accurate compared to the ABM, this deterministic framework is a powerful tool for computing the average behaviour of these cells in this model. Simulating the PDE model on the scaffold in Fig. 5b with a single CPU was around 35 times faster than simulating 2,000 realisations of the ABM.

### 3.3 Impact of micro-rod concentration

The PDE model provides a good approximation of the average prediction from the ABM under scaffolds with varying micro-rod concentrations, as seen in Figs 6a–d (the ABM results represent averages and standard deviations over 500 realisations in each scaffold). Here, we introduce predictions made by the ODE model (Eqs (14)–(16)), which does not consider the spatial distributions of these scaffolds, only their average micro-rod concentration. All results are again scaled up to represent the results expected in the whole culture dish. All models predict that scaffolds with a greater average concentration of micro-rods supports greater overall expansion, which is due to an increased presence of micro-rods to facilitate activation and subsequent proliferation. Scaffolds with more micro-rods introduce more IL-2 to the system as we keep the ratio of loaded IL-2 mass to micro-rod mass, *q*, constant. However, greater activated cell populations in these more concentrated scaffolds cause faster overall IL-2 consumption. As a result, we see similar times for peak T cell expansion across all scaffolds. In [13, 17], T cells are expanded using scaffolds with micro-rod concentrations of 33 µg/mL and 333 µg/mL, and they report that higher micro-rod concentrations are recommended to maximize the number of interactions between T cells and micro-rods, which is supported by our findings.

**Figure 6.**
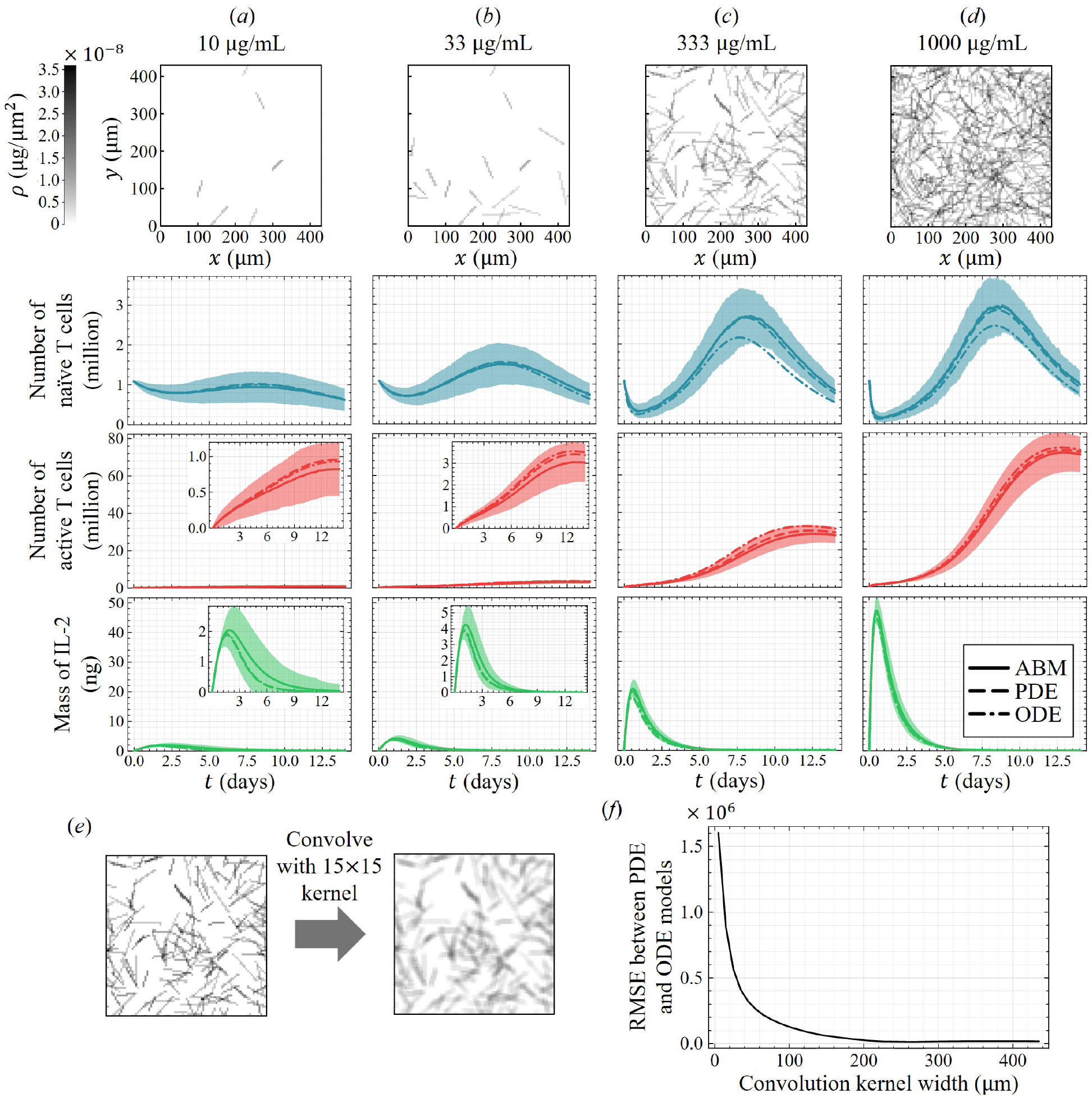
Comparison of T cell population growth and total IL-2 mass between spatial models and the ODE model. *(a-d)* Population growth of naïve T cells (blue), activated T cells (red), and total mass of free IL-2 (green) using the ABM (average and standard deviation of 500 realisations), the PDE model and the ODE model (results scaled up to represent those expected in the whole culture dish) for scaffolds with micro-rod concentrations of *(a)* 10 µg/mL, *(b)* 33 µg/mL, *(c)* 333 µg/mL, and *(d)* 1,000 µg/mL (with zoomed views of activated cell populations and IL-2). *(e,f)* Comparison between the PDE and ODE models for scaffolds convolved with uniform kernels of increasing size, with *(e)* an example of scaffold convolution using a 15 µm × 15 µm kernel and *(f)* the root mean square error (RMSE) between the (scaled up) population predictions of the PDE and ODE models as the convolution kernel increases in size.

The ODE model accurately captures the expansion of activated T cells and the total mass of IL-2, however it is less accurate for predicting naïve T cell populations compared to the spatial models (specifically in Figs 6c,d). Since activated T cells far outnumber naïve T cells at most times, the ODE model also accurately predicts the total number of T cells over time. In Fig. 6f, we compare the ODE model to the PDE model for 333 µg/mL scaffolds in which micro-rod density heterogeneities were smoothed via convolution (for example in Fig. 6e). As the convolution kernel increases in size, the simulated scaffold becomes more well-mixed and, as a result, the population predictions from the PDE model approach that of the ODE model. This assures that, as long as the scaffold is sufficiently well-mixed and similar parameter regimes are implemented, the ODE model may be used to accurately predict the total population of T cells in a computationally efficient manner.

### 3.4 Impact of IL-2 loading and supplementation

Cheung *et al*. report that an important factor in improving T cell expansion over other conventional methods (like with Dynabeads magnetic beads [18]) is the sustained secretion of IL-2 [13]. Experiments suggest that utilising micro-rod secretion nearly doubles T cell expansion compared to experiments in which the same amount of IL-2 is introduced only at the beginning of cell culture [13]. To understand the importance of IL-2 loading in micro-rods we simulated various scenarios involving the secretion of IL-2 from micro-rods (as implemented in all other simulations in this work) and the external supplementation of IL-2 in a representative 333 µg/mL micro-rod scaffold (Fig. 7a). Specifically, we consider cases where IL-2 is not secreted from micro-rods (i.e. 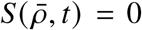) but instead supplemented at the beginning of cell culture or at multiple points throughout co-culture. Here, supplementation is simulated by introducing free IL-2 either at the beginning of co-culture, or at the beginning and at days 2, 4 and 6 (Fig. 7b,c). IL-2 supplementation is a typical technique used when expanding T cells using conventional methods such as with Dynabeads magnetic beads [19–21]. In all cases for our simulations, the total amount of IL-2 introduced to the system is 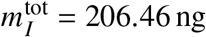, and all results here are scaled up to represent those expected in the whole cell culture (see Sect. 2.5).

**Figure 7.**
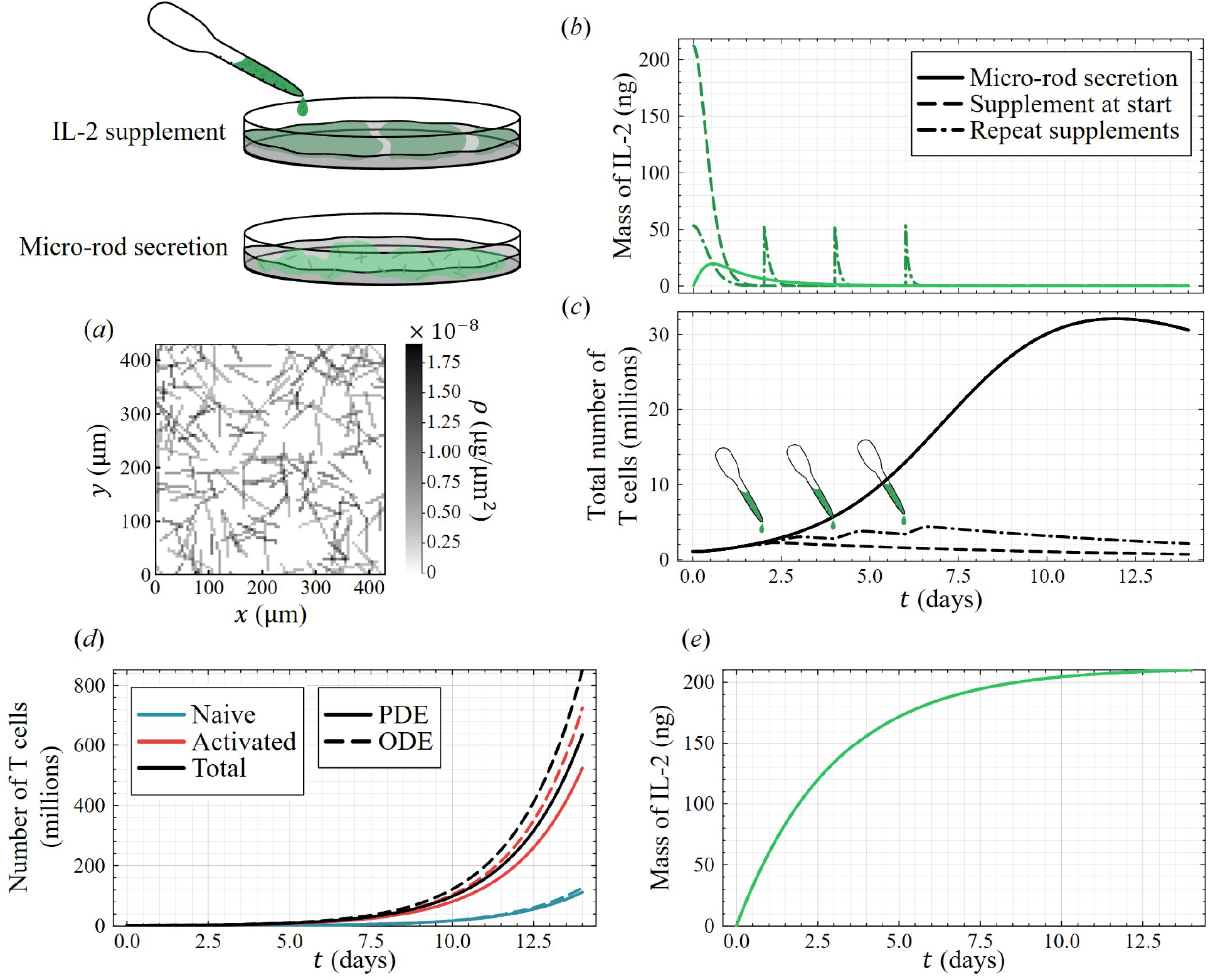
T cell growth response (using the PDE model) when IL-2 is secreted from microrods in a sustained manner and when it is supplemented at the beginning of co-culture or supplemented multiple times throughout co-culture (without micro-rod secretion). *(a)* Microrod mass concentration of a representative 333 µg/mL micro-rod scaffold used to simulate *(b)* the total mass of IL-2 (with consumption by activated T cells) for all scenarios, and *(c)* the corresponding total population of T cells, with diagrams highlighting the IL-2 supplementation at days 2, 4 and 6 (population results scaled up to represent those expected in the whole culture dish). *(d)* T cell populations and *(e)* the total mass of IL-2 (scaled up) in the case where IL-2 is not consumed by activated T cells.

As before, T cells proliferate rapidly until the average IL-2 concentration is around *I*_*p*_ or lower. The sustained release of IL-2 seems to prolong the period over which activated T cells can proliferate rapidly, as the IL-2 stored inside micro-rods is not being consumed by T cells and some IL-2 is always preserved for consumption later during the expansion process. We see similar responses to repeated supplementation where some IL-2 is preserved and dosed later in the cell culture, which slightly prolongs expansion. Consequently, as seen in Fig. 7c, T cell expansion is benefited much more by the sustained release of IL-2 from micro-rods compared to supplementation, and repeatedly supplementing IL-2 promotes slightly greater expansion than only supplementing once at the beginning of cell culture. In this simulation, supplementation only at the beginning of co-culture supports expansion for around 2 days before almost all IL-2 is consumed and the T cell population begins to decline.

In Fig. 7e, we also consider a case where IL-2 is secreted by micro-rods but not consumed by activated T cells, or equivalently, fresh IL-2 is introduced to the system at the same rate that IL-2 is consumed. In this case, Fig. 7d shows that the PDE model predicts unbounded T cell population growth with 600-fold expansion at day 14. This suggests that micro-rod secretion alongside frequent supplementation significantly improves T cell expansion. This seems to be a strategy employed experimentally to ensure that T cells always have access to the necessary IL-2 for proliferation [17]. Additionally, we see that the ODE model predicts faster overall expansion than the PDE model, which suggests that the spatial distribution of micro-rods and T cells is more important for larger scale experiments.

### 3.5 T cell expansion in heterogeneous micro-rod scaffolds

The structure of the micro-rod scaffold is a significant factor for T cell expansion, as these microrods provide the necessary signals for T cell activation. In Fig. 8, we consider the total expansion of T cells in increasingly heterogeneous scaffolds with average micro-rod concentrations of 333 µg/mL (all scaffolds contain the same total amount of loaded IL-2). Each scaffold is generated with different distributions governing the positions of the centres of micro-rods in the *x* and *y* directions. Their heterogeneity, *σ*_*ρ*_, is quantified by the root mean square error between the micro-rod concentration *ρ*(*x, y*) at discrete lattice sites (*i, j*) and the average value of 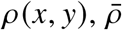:

**Figure 8.**
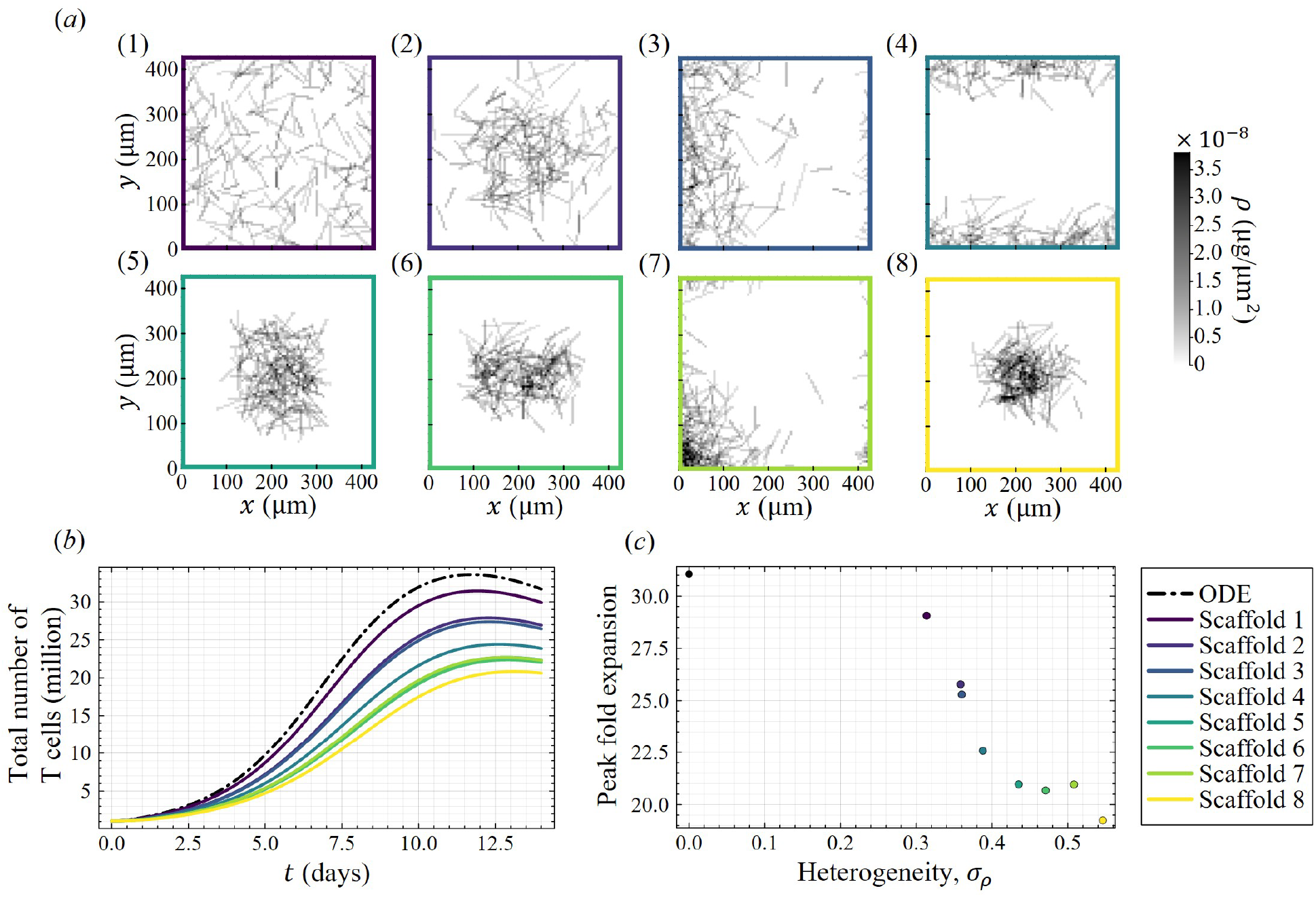
T cell growth within heterogeneous scaffolds. *(a)* Micro-rod mass densities in representative scaffolds (333 µg/mL of micro-rods) of increasing heterogeneity with micro-rod centres (*x, y*) distributed according to (1) (*X,Y*) ~ (*U*(0, *L*),*U*(0, *L*)), (2) (*X,Y*) ~ (*N*(*L*/2, (*L*/5)2), *N*(*L*/2, (*L*/5)2)), (3) (*X,Y*) ~ (Exp(*L*/5),*U*(0, *L*)), (4) (*X,Y*) ~ (*U*(0, *L*), *N*(0, (*L*/10)2)), (5) (*X,Y*) ~ (Triangular(*L*/2, (*L*/4)2), Triangular(*L*/2, (*L*/3)2)), (6) (*X,Y*) ~ (*U*(*L*/4, 3*L*/4), *N*(*L*/2, (*L*/10)2)), (7) (*X,Y*) ~ (Exp(*L*/7), Exp(*L*/7)), and (8) (*X,Y*) ~ (*N*(*L*/2, (*L*/10)2), *N*(*L*/2, (*L*/10)2)). *(b)* Resulting expansion for total T cells (results scaled up to represent those expected in the whole culture dish) and *(c)* the peak fold expansion as scaffolds increase in spatial heterogeneity.

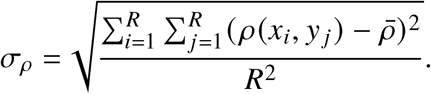

Fig. 8a shows eight representative scaffolds organised in order of increasing heterogeneity, *σ*_*ρ*_. The resulting expansion of total T cells as simulated by the PDE model is shown in Fig. 8b, alongside the prediction made by the ODE model under this average scaffold concentration. Here, and in Fig. 8c, we see that T cell expansion is greatest in scaffolds that are more homogeneous (have smaller *σ*_*ρ*_). The ODE model, which simulates T cell expansion in a completely homogeneous scaffold (*σ*_*ρ*_ = 0 µg/µm^2^), predicts the greatest expansion. In general, as scaffolds become more homogeneous, we see that the maximum fold expansion increases. This is because, in more homogeneous scaffolds, naïve cells are more likely to encounter micro-rods and become activated, as demonstrated in Fig. 4d,e. It is not expected that T cell proliferation is significantly impacted by scaffold heterogeneity, as IL-2 diffuses quickly and the spatial arrangement of micro-rods does not impact the time at which local IL-2 concentrations drop below its critical concentration, *I*_*p*_ (seen by the predictions in Figs 6a–d).

Similarly to the results in Fig. 6f, we see that the ODE model accurately predicts total T cell expansion in the most homogeneous scaffold (lowest value of *σ*_*ρ*_). However, it does not accurately predict the expansion in more hetergeneous scaffolds, which again suggests an importance in spatial modelling, especially in cases where micro-rods are free to move. We do not consider any non-random scaffolds, such as one where the micro-rods are arranged in a grid, as it does not seem practical to do so in experiments due to the fragility of micro-rods [17]. We conducted a similar analysis regarding micro-rod isotropy, and we determined that the orientation of micro-rods does not significantly impact expansion (results not shown).

## 4 Discussion and conclusions

Adoptive T cell therapy (ACT) is an exciting form of immunotherapy where T cells are expanded *ex vivo* to help the immune system fight cancer. Recently, Cheung *et al*. (2018) developed microrod scaffolds that can be used to activate and expand T cells more effectively than conventional methods for ACT. In this work, we presented a hybrid agent-based model of T cell expansion in activating micro-rod scaffolds, in which T cells are modelled stochastically and IL-2 is governed by a T cell-dependent ODE. We computed its continuum limit in the form of a system of differential equations, where T cells are governed by PDEs that describe their diffusion, haptotactic advection, and reactions to micro-rods and IL-2. Finally, we considered a mean-field approximation of the PDE model, which allows us to estimate the average growth of T cell populations.

Each of our three modelling formulations (hybrid ABM, PDE and ODE) provided us with useful insights into the activation and expansion of T cells in micro-rod scaffolds. Simulating with the ABM allowed us to determine that T cell expansion is driven by a positive feedback where individual T cells are activated by micro-rods and so proliferate to produce daughter cells that are subsequently activated by the same micro-rod. The PDE model closely approximated the average result of the ABM, both spatially and in terms of total population, under a wide variety of experimental designs and with greater computational efficiency than the ABM. However, unlike predictions by the PDE model, ABM simulations capture the variability that we may expect during T cell expansion, which more accurately models the variability seen in experiments [13, 17]. The ODE model provided a computationally efficient framework which sufficiently approximates predictions by the PDE model in homogeneous scaffolds, and could easily be used to investigate experimental modifications to the micro-rod platform.

Comparison of the ODE model to the hybrid ABM suggested that the spatial arrangement of micro-rods and T cells is an important factor for understanding and modelling T cell growth. Using a mean-field approximation, we generated a non-spatial system of ODEs, which accurately predicted total T cell expansion and evolution of IL-2 in homogeneous scaffolds. Interestingly, the ODE model inaccurately predicted T cell expansion in heterogeneous scaffolds, as well as in cases where IL-2 remains well-stocked. This suggests that the spatial arrangement of micro-rods is crucial to understanding T cell expansion in micro-rod scaffolds, as this is the component of the system not captured by the ODE model.

Simulation results indicated that the loading of IL-2 into micro-rods benefits expansion as the IL-2 molecules stored inside micro-rods are not consumed during the early stages of expansion. The standard method of simply supplementing IL-2 at the beginning of (or at multiple discrete times throughout) T cell culture, such as experiments in [13, 19], resulted in a faster depletion of IL-2 and a subsequent decline in T cell populations. We also found that combining secretion and supplementation to mitigate or reduce the effects of IL-2 consumption, as is done when expanding T cells with micro-rod scaffolds [13], significantly improves T cell expansion compared to either method individually. In practice, the size of micro-rod pores may be tuned to reduce the rate of secretion and further prolong T cell expansion, or more IL-2 may be loaded into micro-rods to provide activated T cells with more reproductive cues throughout expansion. These results likely follow for any other signaling molecules that may be loaded into the micro-rods, which provides further opportunities for extending these models.

By analysing T cell expansion in heterogeneous scaffolds, we determined a correlation between the distribution of micro-rods and T cell expansion. Expansion was greatest in homogeneous scaffolds as this provided more coverage for T cell activation, whereas activation was less likely to occur in more heterogeneous scaffolds due to the inaccessibility of micro-rods. Real experiments suggest that scaffolds are initially homogeneous and T cells reform scaffolds into heterogeneous clusters of T cells and micro-rods over time [13]. In this case, our results suggested that the the initial homogeneity of the scaffold maximises activation at the beginning of cell culture. Although we do not model the movement of micro-rods, our findings suggested that the subsequent formation of clusters may promote rapid expansion as new naïve cells are produced near micro-rods, which facilitate further activation and proliferation. This is supported by our findings which suggested that T cells activate more quickly in homogeneous scaffolds, but subsequent generations of T cells activate more quickly when near heterogeneous clusters of micro-rods.

### 4.1 Limitations and future work

For simplicity, we chose not to model the effect of gradual scaffold reformation, however, our results suggested that the movement of micro-rods may be important for T cell expansion. In future work, we aim to modify our stochastic framework such that micro-rods adhere to T cells and move due to random T cell motion, and investigate the formation of large cell-rod clusters and their impact on T cell expansion. We expect that this will require the addition of volume filling effects to our model to ensure that T cell concentrations within these clusters do not blow up in finite time due to haptotaxis [27]. A non-spatial model, such as the ODE model presented in this work, is unable to capture the effects of gradual scaffold reformation on T cell expansion, which emphasises the importance of spatial modelling in this system. Due to the flexibility of agent-based modelling, we expect that our stochastic model may easily be modified to predict T cell expansion using other experimental methods. We intend to investigate T cell expansion with other artificial antigen-presenting cells, such as Dynabeads magnetic beads [18].

All models predict that T cell populations expand more quickly in scaffolds with higher average micro-rod concentrations, but there is likely an upper bound for micro-rod concentrations that are feasible in practice. Our current model does not capture volume filling effects, however, we speculate that scaffolds with higher average micro-rod concentration will limit T cell movement and expansion may be reduced as a result. For both micro-rod concentrations implemented experimentally (33 µg/mL and 333 µg/mL), our model predicts T cell expansion lower than what is reported in the experiments by Cheung *et al*. [13]. In practice, more IL-2 is supplemented over time to ensure that T cell proliferation is prolonged [17], and when we increase the amount of IL-2 available, we obtain values closer to that of the experiments.

Our simulation results support the finding that IL-2 secretion improves expansion, but experimental results do not indicate such a drastic difference in expansion between these cases as our simulations suggest [13]. Cheung *et al*. also present results which suggest that increasing the concentrations of micro-rods and activating stimulus decreases the total expansion [13], which is not reflected in our predictions. They do not provide reasons for this observation, so this is perhaps a phenomenon that is not well understood and may be investigated further using these mathematical frameworks. While it may be possible to vary parameters in our model to match their reported T cell expansion, there is currently insufficient data to predict the expansion that they report with certainty. It is possible that our model does not fully capture some relevant biological mechanics involved with T cell proliferation and interactions with micro-rods.

Our model simplifies some biological dynamics of T cell expansion; we do not capture the influence of effector and helper T cells separately, and we assume that micro-rods retain their activating functionalities throughout expansion. In reality, experiments suggest that the ratio of effector-to-helper T cells is dependent on the micro-rod formulation [13, 17], which is an important consideration for ACT. Also, the supported lipid bilayer surrounding micro-rods degrades over time, which may reduce their activating capability [13]. Future iterations of this model will introduce effector and helper T cells, and consider the effects of micro-rod biodegradation on T cell activation.

Finally, we assumed that activation is binary, but this is not the case as T cells can exhibit ranges of activation, which may impact proliferation rates during cell culture [3]. Additionally, T cells may become dysfunctional due to insufficient activation (anergy) or prolonged activation (exhaustion) [47–49]. In a similar sense, T cells may also become dysfunctional due to old age (senescence), which hinders their ability to reproduce [48]. Avoiding T cell dysfunction is a crucial factor for effective applications of ACT [10]. Experiments suggest that T cells expanded using micro-rod scaffolds exhibit a range of dysfunctionality depending on the formulation of micro-rods or scaffold structure [13, 17]. As such, we intend to introduce T cell dysfunction to our model so that we may investigate experimental designs that minimise dysfunction while maximising T cell expansion. Assuming that the ODE model continues to accurately predict T cell populations in homogeneous scaffolds after these model changes, it may be used to efficiently determine non-spatial parameter regimes that optimise T cell expansion.

Our model is one of the first mathematical and computational model of *ex vivo* T cell expansion for ACT. Our hybrid-ABM provides simulations *in silico* at the cellular level, whereas our continuum models provide predictions of the average distributions of T cells and population-level behaviour in micro-rod scaffolds. Our mathematical frameworks will allow for future optimisations of the micro-rod scaffold platform that will hopefully improve the speed and efficiency of T cell expansion for adoptive T cell therapy.

## Acknowledgements

P.R.B. and A.L.J. acknowledge support from the Australian Research Council (ARC) Discovery Project (DP) DP230100025. A.L.J. acknowledges support from the ARC Discovery Early Career Researcher Award (DECRA) DE240100650. M.S.L. acknowledges support from the Australian Government Research Training Program (RTP) Scholarship.

## Appendix A Model fitting

Fig. 9 shows the expansion of T cells under constant concentrations of IL-2 [35], similarly to Fig. 3a. Exponential models are fitted to this data and the resulting growth rates are used for fitting the T cell proliferation rate, *r*_*p*_ (*I*) (see Fig. 3b and Sect. 2.4 for details on the fitted models).

**Figure 9.**
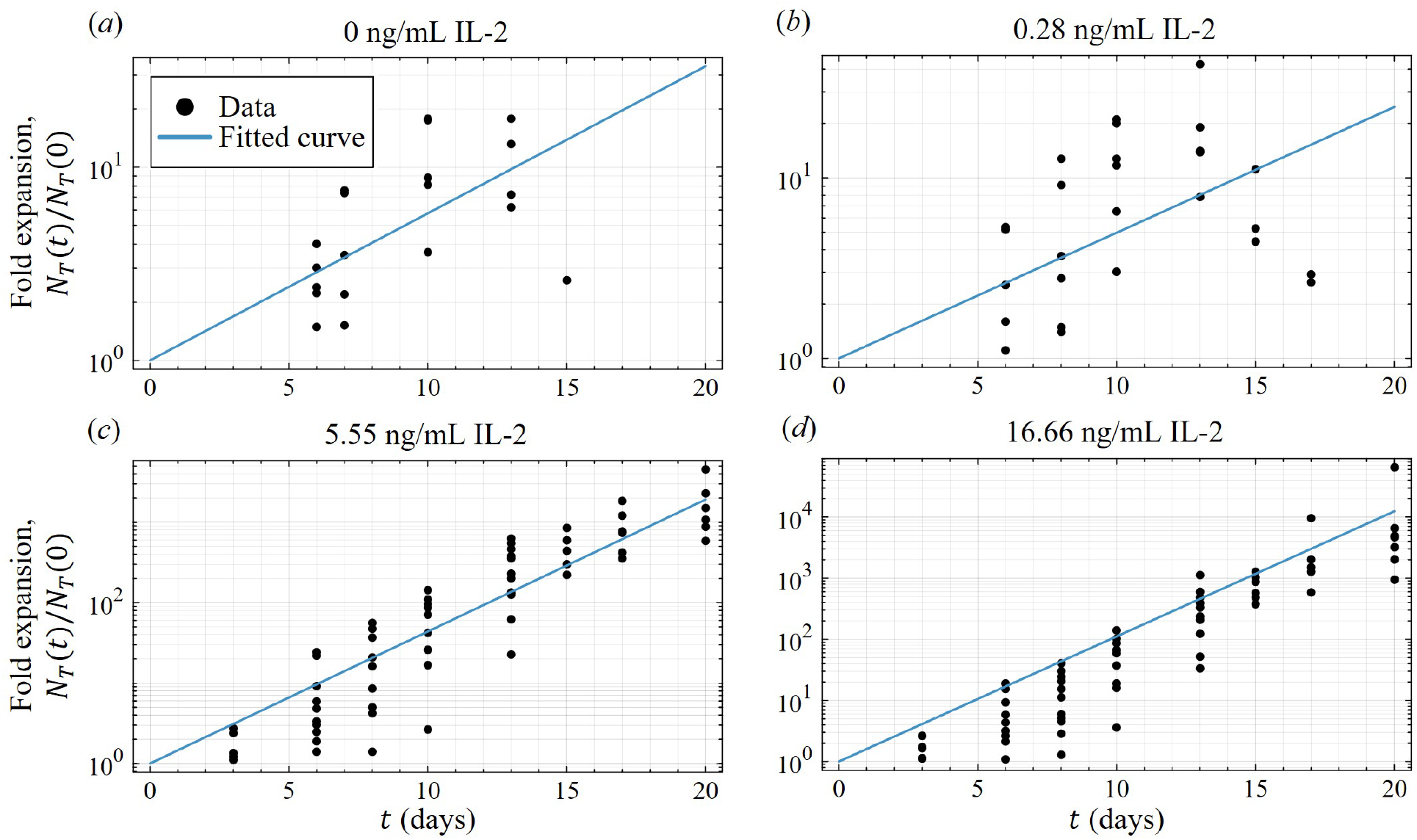
Exponential fits of T cell expansion used to recover proliferation rates under IL-2 concentrations of *(a)* 0 ng/mL, *(b)* 0.28 ng/mL, *(c)* 5.55 ng/mL, and *(d)* 16.66 ng/mL (data reproduced from [35]).

## Appendix B PDE discretisation

The PDEs in Eqs (11)–(12), are solved numerically following spatial discretisation using the FVM and temporal discretisation using the theta method. The square domain is discretised into a grid with vertex-centred nodes (*i, j*) separated by *ε* = 5 µm in both *x* and *y* directions to match the lattice used in the ABM simulation, where again *i, j* = 1, 2, …, *R* and *R* = 86. We may consider each lattice site to be a control volume *Ω*_*i, j*_ surrounding node (*i, j*), which has an area of *ε*^2^. Considering the PDE governing naïve T cells in Eq. (11) for example, we introduce *Q* = −[*r*_*d*_ + *r*_*a*_ (*ρ*)]*n* + *r*_*p*_ (*I*)*a*, and integrate over the control volume:

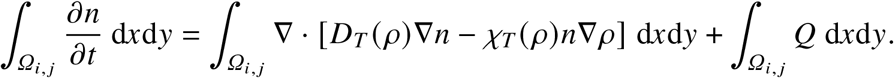

Introducing spatial averages 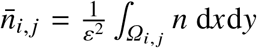 and 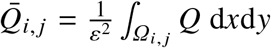, and applying Gauss’ divergence theorem,

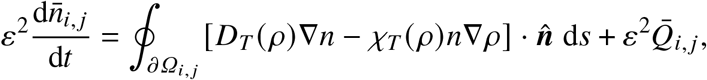

where 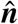 is the outward-facing unit normal on the control volume’s boundary, *∂ Ω*_*i, j*_. Approximate 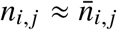 and 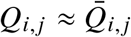 (*Q* evaluated at (*x*_*i*_, *y*_*j*_)), and represent the line integral over *∂ Ω*_*i, j*_ as the sum of contributions over each face of *Ω*_*i, j*_. For interior nodes,

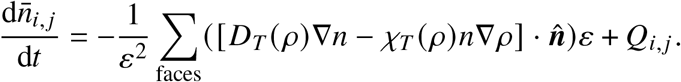

The system of ODEs is arranged into the form 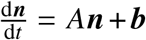, and the Crank–Nicolson method is used to discretise time where time *t* = *k τ*, resulting in

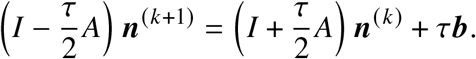

This is implemented and solved iteratively from ***n***^(0)^ until some final simulation time using Julia’s backslash operator, and with use of sparse storage for *A*. Here we use the same timestep as used in the ABM, *τ* = 1 min, for ease of comparison. Under these parameter choices, 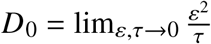 converges to 25 µm^2^/min.

## Notes

### Competing Interest Statement

The authors have declared no competing interest.

